# Plasma proteomics reveals tissue-specific cell death and mediators of cell-cell interactions in severe COVID-19 patients

**DOI:** 10.1101/2020.11.02.365536

**Authors:** Michael R. Filbin, Arnav Mehta, Alexis M. Schneider, Kyle R. Kays, Jamey R. Guess, Matteo Gentili, Bánk G. Fenyves, Nicole C. Charland, Anna L.K. Gonye, Irena Gushterova, Hargun K. Khanna, Thomas J. LaSalle, Kendall M. Lavin-Parsons, Brendan M. Lilly, Carl L. Lodenstein, Kasidet Manakongtreecheep, Justin D. Margolin, Brenna N. McKaig, Maricarmen Rojas-Lopez, Brian C. Russo, Nihaarika Sharma, Jessica Tantivit, Molly F. Thomas, Robert E. Gerszten, Graham S. Heimberg, Paul J. Hoover, David J. Lieb, Brian Lin, Debby Ngo, Karin Pelka, Miguel Reyes, Christopher S. Smillie, Avinash Waghray, Thomas E. Wood, Amanda S. Zajac, Lori L. Jennings, Ida Grundberg, Roby P. Bhattacharyya, Blair Alden Parry, Alexandra-Chloé Villani, Moshe Sade-Feldman, Nir Hacohen, Marcia B. Goldberg

**Author notes:** Denotes co-first authors, listed in alphabetical order (Michael R. Filbin, Arnav Mehta). All individuals contributed equally to sample collection and processing, listed in alphabetical order (Nicole C. Charland, Anna L.K. Gonye, Irena Gushterova, Hargun K. Khanna, Thomas J. LaSalle, Kendall M. Lavin-Parsons, Brendan M. Lilly, Carl L. Lodenstein, Kasidet Manakongtreecheep, Justin D. Margolin, Brenna N. McKaig, Maricarmen Rojas-Lopez, Brian C. Russo, Nihaarika Sharma, Jessica Tantivit, Molly F. Thomas). Denotes co-senior authors (Nir Hacohen, Marcia B. Goldberg).

## Abstract

COVID-19 has caused over 1 million deaths globally, yet the cellular mechanisms underlying severe disease remain poorly understood. By analyzing several thousand plasma proteins in 306 COVID-19 patients and 78 symptomatic controls over serial timepoints using two complementary approaches, we uncover COVID-19 host immune and non-immune proteins not previously linked to this disease. Integration of plasma proteomics with nine published scRNAseq datasets shows that SARS-CoV-2 infection upregulates monocyte/macrophage, plasmablast, and T cell effector proteins. By comparing patients who died to severely ill patients who survived, we identify dynamic immunomodulatory and tissue-associated proteins associated with survival, providing insights into which host responses are beneficial and which are detrimental to survival. We identify intracellular death signatures from specific tissues and cell types, and by associating these with angiotensin converting enzyme 2 (ACE2) expression, we map tissue damage associated with severe disease and propose which damage results from direct viral infection rather than from indirect effects of illness. We find that disease severity in lung tissue is driven by myeloid cell phenotypes and cell-cell interactions with lung epithelial cells and T cells. Based on these results, we propose a model of immune and epithelial cell interactions that drive cell-type specific and tissue-specific damage in severe COVID-19.

## Introduction

As of October 2020, severe acute respiratory syndrome coronavirus-2 (SARS-CoV-2) has caused 34 million cases of coronavirus disease 2019 (COVID-19) and more than 1 million deaths globally (https://covid19.who.int/). The host immune response to SARS-CoV-2 varies considerably^1–4^, leading to clinical manifestations ranging from an asymptomatic carrier state to severe illness, organ dysfunction and death^5^. Immune dysfunction is implicated in the pathophysiology of severe disease, involving aspects of both hyper-immune response (activated inflammatory cascades, cytokine storm, tissue infiltrates, damage) and hypo-immune response (relative lymphopenia, impaired T-cell function, impaired interferon antiviral responses, reduced viral clearance)^5–8^. To date, many studies addressing the host immune response to SARS-CoV-2 are limited by small sample sizes or address narrow sets of immune mediators^2,4,9–13^. In a large cohort of acutely-ill patients presenting to a large, urban, academic Emergency Department (ED), we characterized the host proteomic responses to SARS-CoV-2 using two unbiased plasma proteomic methodologies. This analysis uncovers unique protein signatures associated with COVID-19 disease, severity, aging, tissue destruction, and cellular death that we map to specific cell types in the context of relevant clinical phenotypes, thereby providing insight into underlying disease mechanisms.

## Results

### Evolution of viral response and IFN pathway proteins in the plasma of COVID-19 infected patients

We enrolled 384 unique subjects who presented to our ED with acute respiratory distress suspected or known to be due to COVID-19. Of these, 306 patients were subsequently confirmed to have COVID-19 infection. We classified patients by acuity levels A1-A5 on days 0, 3, 7, and 28, derived from the WHO Ordinal Outcomes Scale^14^ (**Fig, 1A**). For confirmed COVID-19 cases, our primary outcome was the maximal acuity (Acuity_max_) within 28 days of enrollment: A1, death within 28 days (N=42, 14%; **Fig. 1A-B**; **Table S1**); A2, intubation, mechanical ventilation, and survival to 28 days (N=67, 22%); A3, hospitalized and requiring supplemental oxygen (N=133, 43%); A4, hospitalized without requiring supplemental oxygen (N=41, 13%); and A5, discharged directly from the ED without subsequently returning and requiring admission within 28 days (N=23, 8%). A1 and A2 were classified as severe (N=109) and A3-A5 as non-severe (N=197).

**Figure 1.**
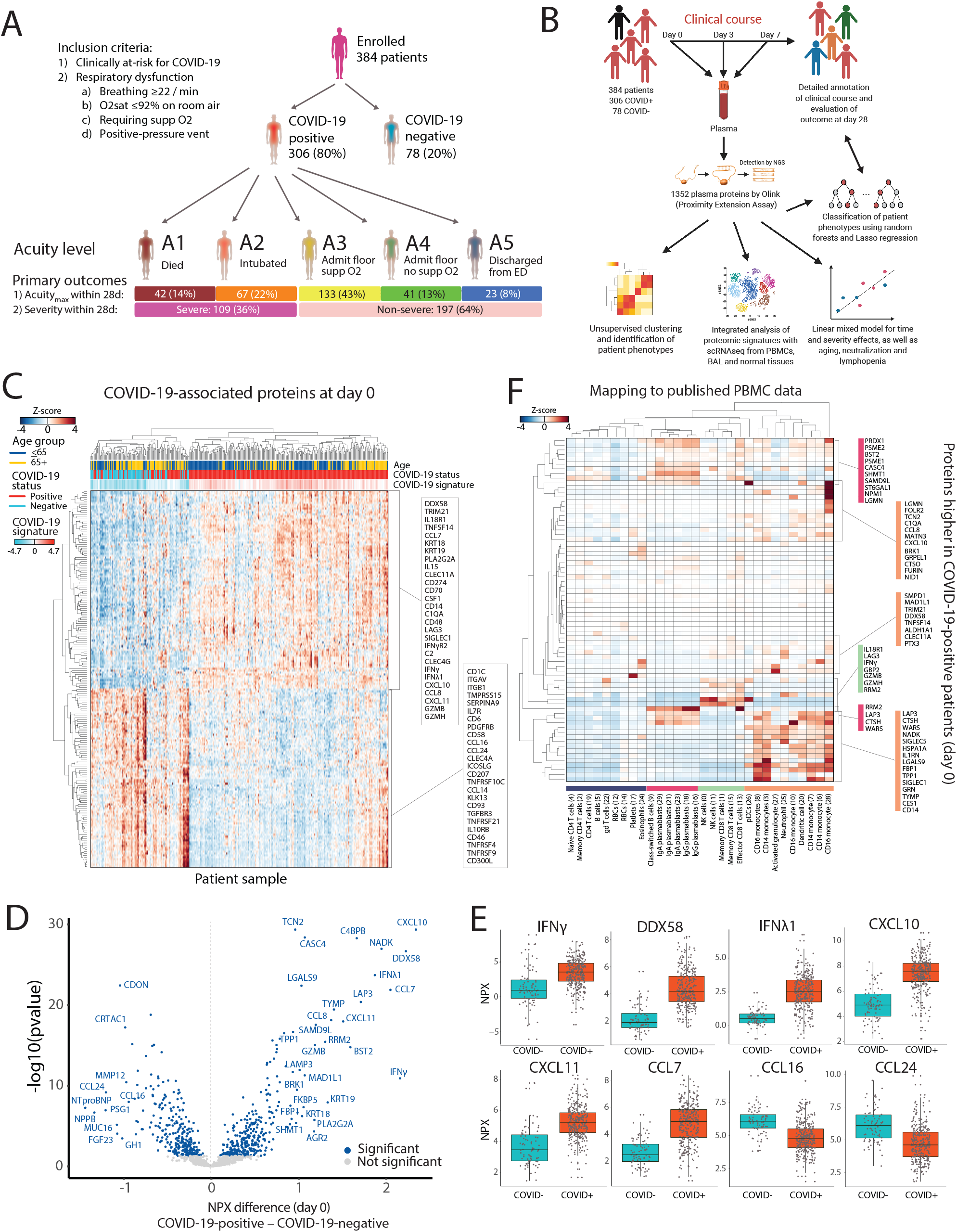
SARS-CoV-2 infection induces viral response and IFN-pathway proteins detected in patient plasma. (A) Schematic of all patients in our study cohort: 306 COVID-19-infected patients and 78 symptomatic COVID-19-negative controls. Inclusion criteria are indicated. Maximal acuity level within 28 days (Acuity_max_) is shown for COVID-19-infected patients (A1, most severe, to A5, least severe), with N, proportion of patients, and severe versus non-severe group assignment. (B) Schematic of study methodology. (C)-(E) Differentially expressed proteins by COVID-19 status. Linear model fitting each Olink protein, with COVID-19 status as a main effect and putative confounders as covariates (see **online methods**). p-values were calculated to account for false discovery rate (FDR) < 0.05, Benjamini-Hochberg method. (C) Heatmap of the top 200 differentially expressed proteins between COVID-19-positive and negative patients. Each row represents expression of an individual protein over the entire cohort; each cell represents the Z score of protein expression for all measurements across a row. COVID-19 signature scores were calculated by taking the mean Z-score of the top 25 differentially expressed proteins in COVID-19-positive patients minus the top 25 differentially expressed proteins in COVID-19-negative patients. (D) Volcano plot showing differentially-expressed proteins based on normalized protein expression (NPX) values between COVID-19-positive and negative patients. Blue circles, significantly differentially-expressed proteins. (E) Box plots of select differentially-expressed viral response and interferon pathway proteins (from panel D), including interferon gamma (IFNγ), DDX58 (or RIG-I), IFNλ1, and the chemokines CXCL10, CXCL11, CCL7, CCL16 and CCL24. (F) Inference of cell of origin by mapping gene expression of differentially-expressed plasma proteins elevated in COVID-19-positive versus negative patients in a scRNAseq peripheral blood cell COVID-19 dataset from Wilk *et al*.^15^. Heatmaps show mean expression of COVID-19-related proteins (y-axis) in immune cell subtypes (x-axis).

COVID-19-positive patients were younger than COVID-19-negative patients (median age 58 vs. 67 years, respectively), with a wide age distribution (**Extended Data Fig. 1)**, and were predominantly Hispanic (54% vs. 15%, respectively). Non-specific inflammatory markers measured clinically, such as C-reactive protein (CRP) and ferritin, were significantly higher in COVID-19-positive versus COVID-19-negative patients; 28-day outcomes were similar between groups (**Extended Data Fig. 1**). Notably, absolute lymphocyte count was not significantly different between groups. We analyzed 1472 unique plasma proteins measured by Proximity Extension Assay (PEA) using the Olink platform (Olink® Explore 1536) (**online methods**) on day 0 (D0; N=383 samples), and for COVID-19-positive patients still hospitalized, also on days 3 (D3, N=218 samples) and 7 (D7, N=136 samples) (**Tables S2, S3**). Unsupervised clustering of D0 samples from 383 patients using protein levels shows clustering by COVID-19 status, age, acuity, ethnicity and kidney disease (**Extended Data Fig. 2A**).

To identify proteins differential between COVID-19-positive and COVID-19-negative patients, we created linear models to fit each of the proteins at D0 with COVID-19 status as a main effect and adjusted for age, demographics, and key comorbidities (**Fig. 1B; Extended Data Figs. 2, 3, Table S4, online methods**). Hierarchical clustering of patients using these differentially-expressed proteins demonstrates a clear separation of the majority of COVID-19-positive from COVID-19-negative patients (**Fig. 1C; Extended Data Fig. 2B**). COVID-19-positive patients displayed higher expression of viral response and interferon pathway proteins, including DDX58 (RIG-I), type II (IFNγ) and type III (IFNλ1) interferons, CCL7, CXCL10, and CXCL11 (**Fig. 1D-E**), with enrichment of proteins in pathways associated with vaccine response, innate immune activation, and T cell function (**Extended Data Fig. 2C-E**). Among chemokines, we found decreased expression of CCL16 and CCL24, among others, in the plasma of COVID-19-positive patients compared to COVID-19-negative patients (**Fig. 1C-E**). Fifty (16%) COVID-19-positive patients clustered with COVID-19-negative patients (**Fig. 1C, left side**) and expressed lower levels of the typical COVID-19 inflammatory signature seen in other COVID-19-positive patients (**Fig. 1C**), yet their mortality was similar to the main cluster of COVID-19-positive patients (**Table S1**); although significantly older than the other COVID-19-positive patients (median age 69 vs 57 years), with more cardiac and kidney comorbidities, this subset is comparably-ill with a distinct low-inflammatory proteomic signature.

To infer immune cell subtype origins of key proteins, we mapped the differential protein expression using the Olink assay in COVID-19-infected patients to single-cell RNA sequencing (scRNAseq) profiles from peripheral blood mononuclear cells (PBMCs) of COVID-19-infected patients^8,15,16^ (**Fig. 1F; Extended Data Fig. 4**). The majority of proteins were selectively expressed in plasmablasts (e.g. RRM2, WARS and PRDX1) and myeloid cells (e.g. CD14, SIGLEC1, SIGLEC10, IL1RN, CCL8 and CXCL10), particularly monocytes and neutrophils (**Fig. 1F; Extended Data Fig. 4**), consistent with the reported expansion and activation of these cell populations during infection^1,15^. Interestingly, monocyte subsets displayed distinct expressions of plasma proteins, with certain subsets demonstrating potent pro-inflammatory responses (**Extended Data Fig. 4**). A smaller group of proteins were expressed strongly in CD8 T cells and NK cells, reflecting cytotoxic responses (IFNγ, GZMB, GZMH and LAG3; **Fig. 1F**).

### Inflammatory pathways associated with severity in COVID-19-positive patients, acute respiratory distress syndrome (ARDS), and their relationship to age

We analyzed protein signatures of COVID-19-positive patients as a function of clinical parameters (**Fig. 1B**). Similar to previous reports^17–19^, Acuity_max_ of COVID-19 patients was significantly correlated with age, D0 acute kidney dysfunction, lactate dehydrogenase (LDH), lymphopenia, acute inflammatory markers (ESR, CRP, D-dimer, and ferritin), and the pre-existing comorbidities kidney disease, diabetes, smoking, and heart disease (**Fig. 2A; Extended Data Fig. 5**). Distinct from some previous reports^17–19^, Acuity_max_ was not significantly correlated with race, ethnicity, or body mass index (BMI) (**Extended Data Fig. 5**). Moreover, neutralization activity (**online methods**) was highly correlated with inflammatory markers and absolute neutrophil count (ANC), but not with Acuity_max_ (**Fig. 2A; Extended Data Figs. 5, 6**).

**Figure 2.**
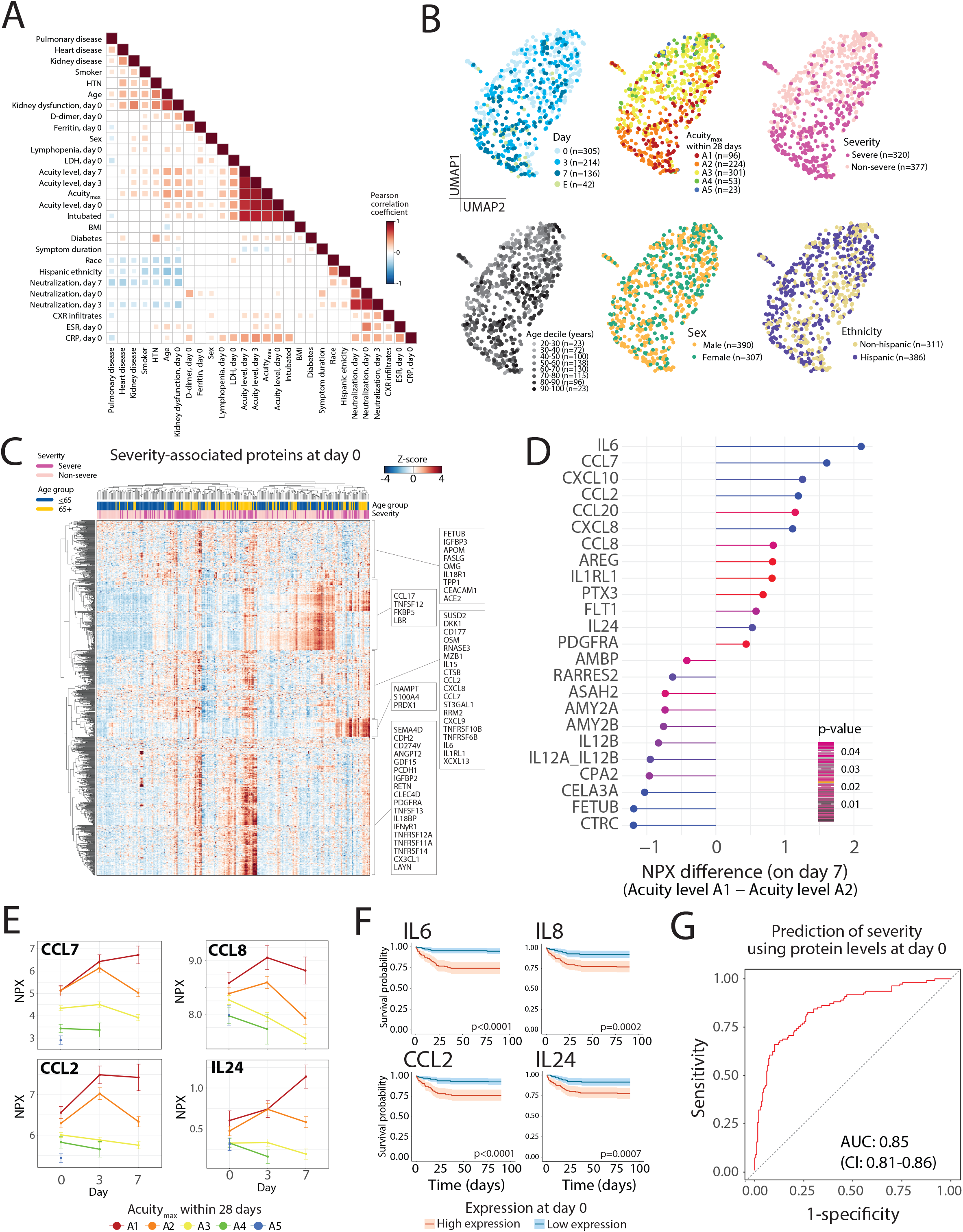
Plasma proteomic biomarkers and predictors of disease severity. (A) Pairwise correlation heatmap of clinically annotated variables for COVID-19-positive patients showing correlations having p<0.05. (B) Unsupervised clustering by UMAP for COVID-19-positive patients at days 0, 3 and 7, color-coded (left to right) by day of sample collection, Acuity_max_ by day 28, severity, age decile, sex, and ethnicity. E, event-driven samples (see **online methods**). (C) Linear mixed model fitting each Olink protein, with severity, timepoint and the interaction of the two terms as main effects and putative confounders as covariates (see **online methods**). Heatmap of significant differentially-expressed proteins between severe and non-severe patients at day 0. Significance of the three model terms determined with an F-test, Satterthwaite degrees of freedom, type III sum of squares. p-values for the three model terms of interest were calculated to account for FDR < 0.05, Benjamini-Hochberg method. Group differences calculated for each significant protein; p-values adjusted using Tukey method. (D)-(F) Linear mixed model fitting each Olink protein, with Acuity_max_, timepoint, and the interaction between the two terms as main effects. Covariates and statistical analysis as in panel (C). (D) Differentially-expressed proteins at day 7 between patients who had Acuity_max_ of A1 (death) versus A2 (ARDS but survived). (E) Point-range plots for select proteins shown in (D). (F) Kaplan-Meier curves for overall survival of patients stratified by higher or lower than median expression of IL-6, IL-8, CCL2, or IL-24. (G) Receiver operating characteristic (ROC) curve showing predictive performance of an elastic net logistic regression classifier of disease severity, for Olink proteins of each patient at day 0. Performance was evaluated using 100 repeats of 5-fold cross validation. Mean area under the curve (AUC) with 95% confidence intervals.

Unsupervised clustering of all samples from COVID-19-positive patients demonstrates that samples cluster by Acuity_max_, severity, age, and time point (**Fig. 2B**). To identify proteins associated with Acuity_max_ levels and severity, we fit linear mixed models (LMMs) to protein values with either Acuity_max_ or severity, and time as main effects, and with covariates age, demographics, and key comorbidities (**Extended Data Figs. 7–9, Table S5-8, online methods**). Because LMMs correct for non-independence in the data, LMMs identify important interactions between variables. At D0, 257 Olink plasma proteins were differentially expressed between severe and non-severe patients, and at D3 and D7, over 700 proteins (**Extended Data Fig. 8B-F, Table S5**). D0 differential proteins were enriched for pathways involved in CD4+ and CD8+ T cell memory responses and IFNα and IL6 signaling in monocytes, among others (**Extended Data Fig. 8E**). Hierarchical clustering of patients by D0 severity-associated proteins revealed distinct clusters of severe patients (**Fig. 2C**), which included a cluster with elevated monocyte inflammatory markers (e.g. OSM, IL6, IL8/CXCL8 and IL15) and a cluster with elevated markers of regeneration and suppressive monocyte proteins (RETN, CLEC4D, GDF15, and PDGFRA). Consistent with this, graphical clustering of patients revealed five patient phenotypes, with three distinct severity patterns starting from D0 (**Extended Data Fig. 10 and Table S9**). As we saw when analyzing proteins by COVID-19-positive status (**Fig. 1**), in the circulation, the majority of proteins associated with severity were most highly transcriptionally expressed in myeloid and plasmablast subsets (**Extended Data Fig. 11**).

Acute respiratory distress syndrome (ARDS) is the leading cause of death in COVID-19 infection. To gain insight into the pathways contributing to death in patients with ARDS, we compared patients who died (A1, N=42, median time to death 9 days [IQR 4-17]) to those receiving mechanical ventilation yet surviving (A2, N=67) (**Tables S6, S7**); by clinical criteria, essentially all patients in these two groups had findings of ARDS. At D7, 24 plasma proteins were significantly differentially expressed between the two groups; amongst those elevated in patients that died were previously reported proinflammatory proteins (IL6^2,4,9,10,20–22^, IL8^2,9,10,21,22^, CXCL10^2,9,10,21^) and previously less well-described monocyte and regeneration proteins, including CCL2, CCL7, CCL8, CCL20, AREG, IL1RL1, PTX3, and FLT1, as well as IL24 (**Fig. 2D,E; Extended Data Fig. 9A,B**). Most of these proinflammatory proteins showed similar elevation and upward trajectories for groups A1 and A2 through D3, followed at D7 by a decline in survivors and a sustained elevation in those who died (**Fig. 2E; Extended Data Fig. 9B**). Plasma levels of these proteins at D0 were associated with poorer survival (**Fig. 2F; Extended Data Fig. 9C**). Interestingly, several exocrine pancreas proteases and protease inhibitors (CTRC, CELA3A, CPA2, CTRB1, AMY2A, AMY2B) were reduced in the plasma of those who died (**Fig. 2D; Extended Data Fig. 9B**); whereas the implication of these in COVID-19 remains uncertain, many have anti-inflammatory effects in mouse models^23–26^. Of note, few patients received the anti-inflammatory dexamethasone, which, since our study, has been established as standard-of-care. Taken together, these findings suggest that survival from COVID-19 ARDS is associated with decreased pro-inflammatory and increased anti-inflammatory responses over time.

### Plasma proteomic signatures predict severity and uncover pathways associated with immune aging

To test whether D0 plasma proteins are predictive of severe disease at later time points, we built a classifier based on the Olink data of severe disease (defined as Acuity_max_ of A1 or A2) using elastic-net logistic regression with cross-validation; D0 plasma proteins levels yielded good predictive performance (AUC 0.85, confidence interval (CI) 0.81-0.86) (**Fig. 2G; Extended Data Fig. 9D, online methods**). Consistent with our LMM results (**FIg. 2C; Table S5**), IL6, IL1RL1, PTX3, and IL1RN were amongst the strongest weighted proteins in the predictor; also amongst top weighted proteins were KRT19, a marker of epithelial damage^27^, and TRIAP1, an inhibitor of apoptosis^28^ (**Extended Data Fig. 9E**). These proteins may therefore serve as predictive biomarkers, which would need to be validated in larger independent cohorts.

As has been previously reported by others, severity was strongly associated with age (**Fig. 2A**). To identify protein pathways specifically associated with age, we looked for proteins with the strongest pairwise correlation with age (**Table 10**) and fit each to an LMM with age and time point as main effects (**Extended Data Fig. 12A, Table S11, online methods**). This approach revealed several severity-associated proteins and proteins previously reported as age-associated^29–31^ (GFAP, EGFR, MLN, EDA2R) that separated patients by age. Plasma protein expression at D0, D3, or D7 accurately classifies patients by age group using regularized logistic regression (AUC 0.93, 0.90 and 0.86, respectively) and identifies several age-related proteins independent of (IL2RB, GZMA, FLT3, CD27, IL17D) or associated with (RSPO3, NT-proBNP, CHI3L1, CD59, LAYN) disease severity (**Extended Data Fig. 13**). To identify temporal protein patterns indicative of immune aging, we fit each protein to a LMM with disease severity, time point, and age (≤ or > 65 years) as main effects (**Table S12, online methods**), identifying four distinct dominant expression patterns between young and aged, and non-severe and severe patients (**Extended Data Fig. 12B**), that together highlight unique age-related responses to COVID-19 infection.

### Plasma protein signatures of tissue damage highlight organ toxicity in severe COVID-19 infection

To elucidate patterns of tissue damage, we derived tissue-specific plasma proteomic signatures by analyzing samples using the SomaScan platform^32^, as this provides complementary sampling of a broad range of the plasma proteome, detecting over 4400 proteins via an aptamer-based technology. LMMs of severity and Acuity_max_ resulted in largely overlapping sets of differentially expressed proteins with the Olink platform (hypergeometric test p=0.002, 69% of severity-associated proteins overlapped with those identified by Olink data) (**Table S13, online methods**). To derive organ-specific expression signatures, we leveraged gene expression data from the Genotype-Tissue Expression (GTEx)^33^ portal (**online methods**) and verified the cell type of origin of each protein in a given signature within each tissue of interest, reassuringly finding that our derived signatures were expressed primarily in epithelial cells (**Extended Data Fig. 14**). We then identified plasma proteins in our datasets that overlapped with these gene sets (**Table S14**), looking specifically at proteins that are intracellular and thus are likely to leak into circulation as a result of tissue damage (**Fig. 3A; Extended Data Fig. 15A**). When possible, we validated our organ-specific signatures (**Table S14**) against lab values measured clinically; this showed significant correlations of our heart signature with clinical troponin levels (R=0.52, p < 0.001), our liver signature with clinical ALT (R=0.6, p < 0.001) and AST (R=0.46, p < 0.001), our skeletal muscle signature with clinical CPK (R=0.6, p < 0.001), and negative correlations between our kidney signature and GFR (R=-0.36, p < 0.001) (**Fig. 3B**; **Extended Data Fig. 16)**.

**Figure 3.**
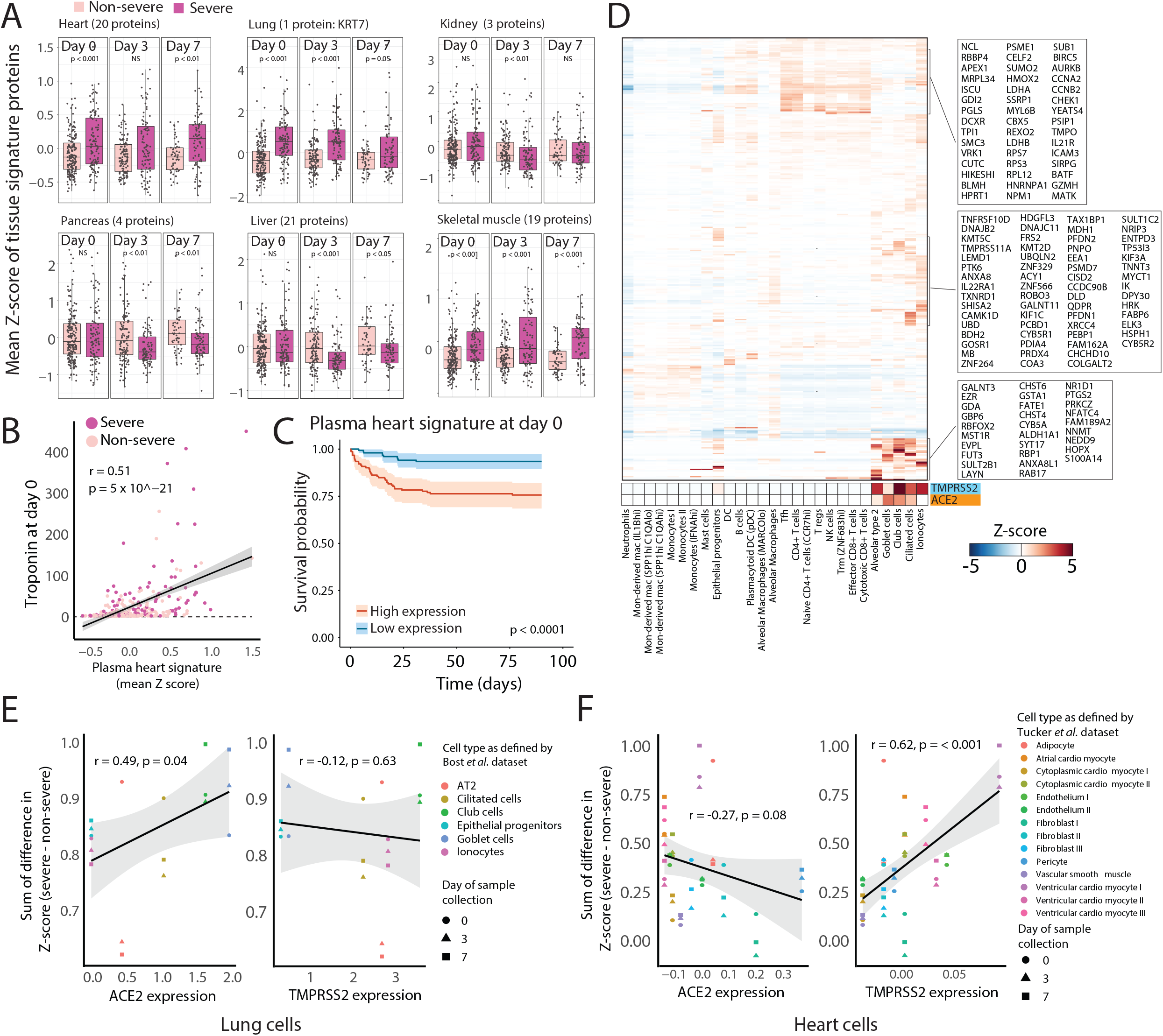
Severe COVID-19-positive patients display elevated plasma markers of cell death from heart, lung, and skeletal muscle. (A) Expression of tissue-specific plasma protein signatures in severe versus non-severe patients at each timepoint. (B) Scatter plot showing the correlation of the D0 plasma heart signature as derived in (A) with D0 clinical troponin measurements. (C) Kaplan-Meier curve showing overall survival of patients with high or low expression (above or below median expression level) of the derived plasma heart signature in (A). (D) Heatmap showing mean gene expression per cell type of severity-associated intracellular plasma proteins at D0 derived from SomaScan data that map to scRNAseq of BAL fluid^34^, with TMPRSS2 and ACE2 expression indicated. (E)-(F) Scatter plots showing the difference between severe and non-severe patients of lung (E) and heart (F) cell-specific intracellular death scores, derived from expression of differentially-expressed proteins at each timepoint versus cell type-specific ACE2 and TMPRSS2 expression levels from scRNAseq of BAL fluid^34^ (E) or heart single-nucleus RNA-seq data^53^ (F).

In patients with severe COVID-19 (A1, A2), heart, lung, and skeletal muscle intracellular protein signatures were elevated in plasma as early as D0 and this elevation persisted to D7 (**Fig. 3A**). Consistent with this, elevated heart and skeletal muscle protein signatures at D0 portended poor overall survival, but this was not the case for lung, kidney, liver, and pancreas signatures (**Fig. 3C; Table S15**). Interestingly, in severely ill patients, liver and pancreas intracellular protein signatures were decreased in the plasma at D3 and D7 (**Fig. 3A**); over the same timeframe, liver-specific secreted proteins were decreased, whereas pancreas-specific secreted proteins were increased (**Extended Data Fig. 17**). Together these results suggest that COVID-19 illness drives damage to the heart and lung and that this damage signature can be detected early in the circulation upon hospital presentation.

### Lung damage in severe COVID-19 is a result of cell death within epithelial cells

We mapped intracellular severity-associated plasma proteins to specific tissues and cell types using scRNAseq datasets (**Extended Data Figs. 18, 19, online methods**). Analysis of scRNAseq datasets from healthy tissues revealed that plasma proteins at D0 (**Table S5**) are expressed in macrophage subsets and epithelial cells, with notably increased relative expression in proximal tubule cells in the kidney, hepatocytes in the liver, and stellate, ductal, and acinar cells in the pancreas (**Extended Data Fig. 19**). By analyzing expression of intracellular severity-associated proteins (**Table S5**) in the more physiologically-relevant context of single cells of bronchoalveolar lavage (BAL) fluid from COVID-19 infected patients^34^, we found distinct clusters of proteins expressed within lung epithelial cells and within T cells, with little expression in tissue-associated myeloid or B cells (**Fig. 3D**). These tissue-associated patterns highlight the distinct contributions to the plasma proteome of cell types in the circulation and those in tissue (**Fig. 3D; Extended Data Fig. 11**). Within lung epithelial cells, expression of severity-associated proteins correlated with ACE2 expression (R=0.49, p = 0.04), but not with TMPRSS2 expression (**Fig. 3F; Extended Data Fig. 15B**), which suggests that the increased levels of these proteins in plasma results from SARS-CoV-2 infection and death of these cells and that other proteases may be redundant with TMPRSS2 for priming during viral entry. In contrast, heart cell-type plasma signatures did not correlate with ACE2 expression (**Extended Data Fig. 18B**), suggesting that heart damage is largely an indirect effect of the disease process, putatively due to the inflammatory response. The implications of the observed correlation with TMPRSS2 expression on cells (R=0.62, p < 0.001; **Fig. 3F**) is unclear. Similarly, intracellular plasma signatures from cell subsets that do not express ACE2 or TMPRSS2 (**Fig. 3D**) likely results from bystander cell death. Of note, unlike in the blood, a larger subset of severity-associated plasma proteins is expressed in effector and cytotoxic CD8+ T cells and NK cells located within the lung (**Fig. 3D**). Together, our data suggest that death of lung epithelial cells may be a key driver of the pathology of severe disease, and this may be detected from signals in the circulating proteome.

### Cellular communication between lung epithelial cells, T cells, and myeloid cells is associated with disease severity

To gain insights into inflammatory pathways activated in severe disease and given the likely role in disease of elevated levels of circulating proteins, we looked for enrichment of these pathways among plasma proteins that are normally secreted or membrane-bound. Among the D0 severity-associated proteome, NF-kB signaling, NK cell activation, IL6 and IL10 signaling, and Th1 and Th2 activation were significantly enriched (**Extended Data Fig. 17A**). To identify cellular mechanisms regulated by this subset of proteins, we analyzed ligand-receptor interactions^35,36^ using the BAL fluid cell dataset from COVID-19-infected patients^34^ (**Fig. 4A; online methods**). From D0 to D3, there was a dramatic increase in the number of predicted ligand-receptor interactions (**Fig. 4A**), predominantly represented by ligand-receptor interactions occurring in lung epithelial cells, T cells, and mast cells (**Fig. 4B**). Most dramatic were changes in mast cells, driven by their interactions with other mast cells, CD4+ and CD8+ T cell subsets, and epithelial progenitors (**Fig. 4B**). Consistent with this, the mast cell function marker tryptase was differentially expressed between severe and non-severe patients and over time (**Extended Data Fig. 20**).

**Figure 4.**
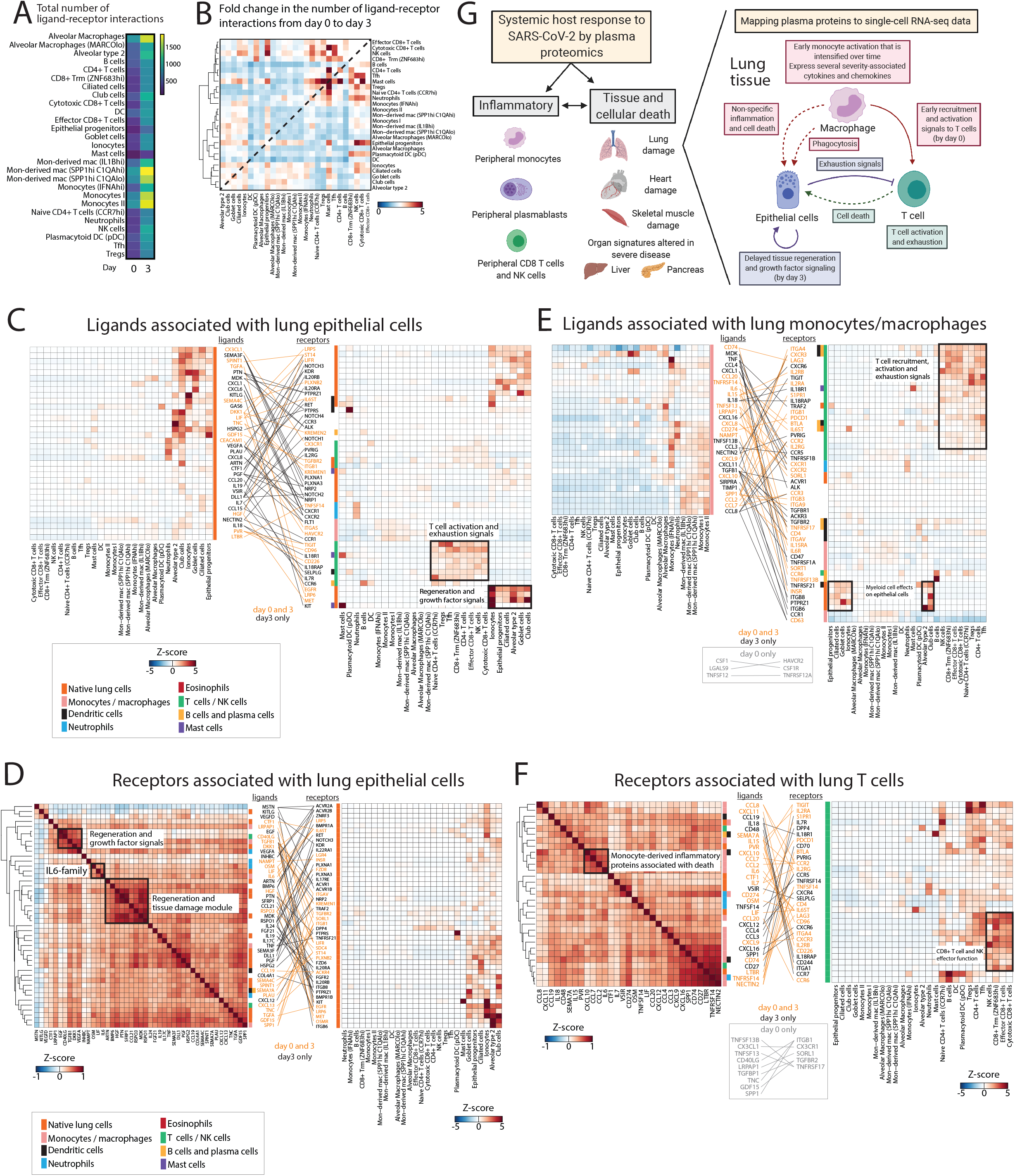
Interactions among lung epithelial cells, monocytes, and T cells drive disease severity and tissue damage. (A) Heatmap showing the total number of ligand-receptor interactions at D0 and D3 inferred from BAL fluid scRNAseq data^34^ using only ligands differentially expressed in the plasma of severe versus non-severe COVID-19-positive patients. (B) Heatmap showing fold-change from D0 to D3 in the number of ligand-receptor interactions between each cell type identified from BAL fluid scRNAseq data^34^. (C) Ligand-receptor contact map between ligands expressed by lung epithelial cells per BAL fluid scRNAseq data^34^ (left) and the respective receptors for these ligands and their cell-specific expression from the same BAL dataset (right). (D) Ligand-receptor contact map between receptors expressed on lung epithelial cells in BAL fluid^34^ (right) and their respective severity-associated plasma ligands from our data (left). Ligand-receptor pairs are those for which the ligand was significantly associated with severity in our models. (E) Ligand-receptor contact map between ligands expressed on monocytes/macrophages in BAL fluid scRNAseq data^34^ (left) and the respective receptors for these ligands and their cell-specific expression from the same BAL dataset (right). (F) As in panel E, but on monocytes/macrophages in BAL fluid. In panels (C)-(F), each cell in the heatmaps represents expression of the listed ligand or protein relative to its expression across all cell types. Ligands and receptors are color-coded (vertical color bar) based on the cell type that demonstrates their highest expression. Ligand-receptor pairs and their connecting lines are color-coded based on the time point (D3 only, or both D0 and D3) for which the interaction was present. (G) Model highlighting contributions to the plasma proteome from circulating immune cells (primarily monocytes, plasmablasts, and CD8+ T and NK cells) and damaged tissues. Temporally-ordered interaction network between monocyte/macrophages, T cells, and lung epithelial cells that drives disease severity.

Whereas lung epithelial cells showed predicted ligand-receptor interactions with other epithelial cells, monocytes and macrophages, and CD8+ T cells, and NK cells, pronounced COVID-19 severity and temporal changes were observed in proteins mediating interactions between epithelial progenitors and cytotoxic/effector CD8+ T cells or NK cells (**Fig. 4B-F**). To better understand the specific pathways mediating disease severity, we constructed mappings of key ligand-receptor relationships of cells in BAL fluid with plasma severity-associated ligands at D0 and D3 (**Extended Data Fig. 21**). Pairings of ligands from lung epithelial cells with receptors on other lung epithelial cells were involved in alveolar maintenance and protection, growth factor signaling, and tissue regeneration (HGF-MET, TGFA-EGFR, DKK1-LRP6, KITLG-KIT, semaphorin-PLXNA receptors, among others; **Fig. 4C**). Moreover, several T cell activating and exhaustion signals originated from lung epithelial cells, mediated by PVR signaling through TIGIT and CD96 as early as D0, and eventual activation of IL7 and IL18 pathways at D3 (**Fig. 4C**). To better understand the pathways affecting lung epithelial cells, we focused on the receptors expressed in these cells, specifically those that have ligands that were severity-associated in our model (**Fig. 4D**). A correlation matrix using plasma expression levels of these ligands identified coregulated groups of proteins that act on lung epithelial cells. Amongst these were clear protein modules for regeneration and growth factor signaling (module 1: EGF, TGFB1, DKK1, VEGFA; module 2: BMP6, HGF, RSPO3, RSPO1) and for IL6 pathway signaling (OSM, LIF, IL6) (**Fig. 4D**).

Amongst the severity-associated ligands expressed on monocytes/macrophages are many known to be associated with death in COVID-19 (IL8, CXCL8, CCL2, CCL7, CCL8, and CCL20; **Figs. 2D, 4E**). We found several previously poorly characterized monocyte/macrophage ligand-receptor interactions with lung epithelial cells and other myeloid cells, most apparent at D3, that may be involved in driving later-stage damage and regulating phagocytosis (e.g. TGFB1-ITGB6 and ITGB8, SPP1-ITGAV, SIRP1a-CD47; **Fig. 4E**). The interaction of TGFB1 with its receptor ITGB6/8 likely suppresses beneficial immune responses, as it releases active TGFB1 from its inhibitor latency-associated peptide^38,39^. Active TGFB1 suppresses cytotoxic T cells, potentially inhibiting clearance of SARS-CoV-2-infected cells, inhibits proliferation of naïve T cells and induces growth arrest of B cells, potentially contributing to COVID-19-associated lymphopenia, and promotes differentiation of peripheral Tregs^40^. Of note, the majority of monocyte ligands function in T cell recruitment, activation, and exhaustion as early as D0 (e.g., CXCL9-CXCR3, CXCL10-CXCR3, IL15-IL2R, CD74-LAG3, CD274-PDCD1; **Fig. 4E**). Consistent with this, amongst the subset of severity-associated monocyte ligands that affect receptors on T cells is a coexpressed module of monocyte death-associated cytokines/chemokines (CXCL10, CCL7, CCL8, CCL2; **Fig. 4F**). Also notable are two coexpressed modules of receptors on T cells associated with T cell activation and exhaustion (IL18, IL15, and PVR, and CXCL16, CD74 and CD27), the ligands for which are expressed on monocytes (**Fig. 4F**). Based on these data, we propose a model of COVID-19-induced immune and cellular responses within the lower airways in which early monocyte activation drives T cell recruitment, activation, and exhaustion, leading to delayed activation of additional monocyte inflammatory pathways and repair and regeneration within lung epithelial cells (**Fig. 4G**).

## Discussion

This analysis of the plasma proteomes of 384 acutely-ill patients with COVID-19 and non-COVID-19 illnesses provides a comprehensive longitudinal summary of the systemic host response to SARS-CoV-2. We found that COVID-19 patients have dramatically different plasma proteomic profiles than COVID-19-negative controls with respiratory distress (**Fig. 1**). We broadly characterize over 600 plasma proteins implicated in the host viral and inflammatory response to SARS-CoV-2. Amongst COVID-19-infected patients, a small subset had inflammatory signatures similar to COVID-19-negative controls but had outcomes similar to those seen with other COVID-19-positive patients. Over 250 proteins were independently associated with COVID-19 severity, with several inflammatory mediators associated with death in ARDS patients, including previously identified markers (IL6^2,4,9,10,20–22^, IL8^2,9,10,21,22^ and CXCL10^2,9,10,21^) and several novel markers (CCL2, CCL7, CCL8, CCL20, AREG, IL1RL1, FLT1 and IL24) (**Fig. 2**). Importantly, plasma proteins at D0 are strongly predictive of disease severity. These may identify pathways associated with severity that are amenable to existing or new therapeutics. Furthermore, they can be incorporated into diagnostics that stratify high-risk patients for tailored therapies or earlier intervention.

By leveraging scRNAseq datasets from PBMCs of COVID-19 patients and healthy tissues, we deconvoluted the relative contribution of different compartments to the plasma proteome, finding a subset of severity-associated plasma proteins expressed in circulating monocytes and plasmablasts, and a smaller subset in circulating T cells and NK cells. To understand the effect of COVID-19 on host tissues, we leveraged gene expression data from GTEX to derive organ-specific cellular death signatures. By applying the resulting unbiased set of intracellular proteins to severity-associated proteins in plasma proteomes, we demonstrate that severe patients have early signals of heart, lung, and skeletal muscle tissue damage (**Fig. 3**). Our derived protein signatures correlate with clinical metrics of tissue-specific cellular damage and, by using scRNAseq data, primarily show gene expression in epithelial cells within the respective tissues, providing support to their validity. Our plasma organ-specific death signatures will have broader utility as a potential liquid biopsy for organ damage and can be used to interpret the plasma proteome in the context of tissue-specific cell death and inflammation.

By applying expression profiling of cells from BAL fluid to our intracellular cell death signatures, we demonstrate that the severity-associated proteome is significantly associated with cell-type specific ACE2 gene expression, suggesting that direct infection of lung epithelial cells leads to cell death that is measurable in plasma (**Fig. 3**). Of note, we make the assumption that circulating factors reflect the effect of secreted factors in lung tissue, an assumption that requires validation in future studies. By analyzing the interactions of circulating ligands with receptors within cells in BAL fluid, we discover a temporal order of cellular communication in the lung that drives disease severity (**Fig. 4**). This starts with early activation of monocytes/macrophages and T cell recruitment into tissues alongside activation and early expression of exhaustion markers. Death of lung epithelial cells in turn activates regenerative and growth factor signaling, and phagocytosis of cells by macrophages (**Fig. 4G**), consistent with proposed models of altered differentiation dynamics of nasal epithelial cells upon SARS-CoV-2 infection^37^. By discriminating specific cell types likely dying from direct viral infection from those likely dying from indirect effects of the illness, we provide new details on the pathogenesis of the disease.

In conclusion, we studied several thousand circulating proteins within the plasma proteome in a large number of symptomatic COVID-19 patients and acutely-ill non-COVID-19 controls upon hospital arrival and during the first week of hospitalization. We performed adjusted analyses to determine plasma proteins that are independently-associated with COVID-19 disease and illness severity. In patients with ARDS, we identified inflammation-related proteins associated with death, and non-inflammatory, potentially protective proteins that are upregulated in those who survive. By mapping differentially-expressed proteins to published scRNAseq data, we infer cellular origins of relevant circulating proteins and provide an important link between cellular immune responses and tissue damage in severe disease to the clinically-accessible plasma proteome. This work lays the foundation for understanding how the plasma proteome can be leveraged to gain insights into underlying disease pathways and potential therapeutic targets, and to develop diagnostics that can be used to stratify high-risk patients as candidates for tailored therapies and earlier interventions.

## Methods

### Patient cohort and clinical data collection

Patients were enrolled in the Emergency Department (ED) of a large, urban, academic hospital from 3/24/2020 to 4/30/2020 in Boston during the peak of the COVID-19 surge, with an institutional IRB-approved waiver of informed consent. Included were patients 18 years or older with a clinical concern upon ED arrival for COVID-19 and with acute respiratory distress, with at least one of the following: 1) tachypnea (≥22 breaths per minute), 2) oxygen saturation ≤92% on room air, 3) a requirement for supplemental oxygen, or 4) positive-pressure ventilation. The day 0 blood sample (N=384) was obtained concurrent with the initial clinical blood draw in the ED, and day 3 (N=217) and day 7 (N=144) samples were obtained for COVID-19-positive patients, if still hospitalized at those times, yielding 735 samples. In addition, blood was collected from some patients at the time of substantial clinical deterioration (44 samples); these event-driven samples were excluded from linear models. Clinical course was followed to 28 days post-enrollment or until hospital discharge, if that occurred after 28 days. Of all 384 enrolled, 78 (20%) tested negative for SARS-CoV-2; among these, for 50 (64%), suspicion for COVID-19 was very low based on careful retrospective chart review by MRF and RPB, an emergency physician and infectious diseases physician, respectively. Among the remaining 28 patients, COVID-19 was a diagnostic possibility yet most had multiple negative PCR tests throughout their hospital course. These 78 subjects were categorized as controls. We dichotomized COVID-19 subjects by illness severity and outcome into severe (A1-A2) and less severe (A3-A5) groups. Of the 42 COVID-19 patients who died, 24 (57%) received mechanical ventilation and 18 (43%) did not. The latter group was significantly older, many with advanced directives to withhold aggressive care. Demographic, past medical history and clinical data were collected and summarized for each outcome group, using medians with interquartile ranges and proportions with 95% confidence intervals, where appropriate.

### Plasma collection and processing

Blood samples were collected in EDTA tubes and processed no more than 3 hours post blood draw in a Biosafety Level 2+ laboratory on site. Whole blood was diluted with room temperature RPMI medium in a 1:2 ratio to facilitate cell separation for other analyses using the SepMate PBMC isolation tubes (STEMCELL) containing 16 mL Ficoll (GE Healthcare). Diluted whole blood was centrifuged at 1200 *g* for 20 minutes at 20 °C. After centrifugation, plasma (5 mL) was pipetted into 15 mL conical tubes and placed on ice during PBMC separation procedures, centrifuged at 1000 *g* for 5 min at 4°C, aliquoted into cryovials, and stored at −80°C. Study samples (45 μL) were randomly allocated onto 96-well plates based on disease outcome grouping and were treated with 1% Triton X-100 for virus inactivation at room temperature for 2 hrs.

### Olink plasma proteomic assays

The Olink Proximity Extension Assay (PEA) is a technology developed for high-multiplex analysis of proteins using 1 μL of sample. In PEA, oligonucleotide-labelled monoclonal or polyclonal antibodies (PEA probes) are used to bind target proteins in a pair-wise manner thereby preventing all cross-reactive events. Upon binding, the oligonucleotides come in close proximity and hybridize followed by extension generating a unique sequence used for digital identification of the specific protein assay. With recent developments, PEA enables an increased number of 384 multiplex assays and higher throughput using next-generation sequencing (NGS) as a readout method. PEA probe design is based on addition of Illumina adapter sequences, unique barcodes for protein identification and indexes to distinguish samples in multiplex sequencing. The protocol has also been miniaturized and automated using liquid handlers to further improve robustness and maximize output.

The full library (Olink® Explore 1536) consists of 1472 proteins and 48 controls assays divided into four 384-plex panels focused on inflammation, oncology, cardiometabolic and neurology proteins. In each of the four 384-plex panels, overlapping assays of IL-6, IL-8 (CXCL8), and TNF are included for quality control (QC) purposes. Library content is based on target selection of low-abundant inflammation proteins, actively secreted proteins, organ-specific proteins leaked into circulation, drug targets (established and from ongoing clinical trials), and proteins detected in blood by mass spectrometry. Selection, classification, and categorization of proteins were based on using various databases (e.g. Gene Ontology), the Blood Atlas – the human secretome (www.proteinatlas.org), a collaboration with the Institute of Systems Biology, Seattle WA, for tissue-specific proteins, www.clinicaltrials.gov for mapping of drug targets, detection of proteins in blood measured by mass spectrometry and finally, various text-mining approaches identifying protein biomarkers described in the literature. The analytical performance of PEA is carefully validated for each protein assay; performance data are available at www.olink.com. Technical criteria include assessing sensitivity, dynamic range, specificity, precision, scalability, endogenous interference, and detectability in healthy and pathological plasma and serum samples.

In the immune reaction, 2.8 μL of sample is mixed with PEA probes and incubated overnight at 4 °C. Then, a combined extension and pre-amplification mix is added to the incubated samples at room temperature for PCR. The PCR products are pooled before a second PCR step following addition of individual sample index sequences. All samples are thereafter pooled, followed by bead purification and QC of the generated libraries on a Bioanalyzer. Finally, sequencing is performed on a NovaSeq 6000 system using two S1 flow cells with 2 × 50 base read lengths. Counts of known sequences are thereafter translated into normalized protein expression (NPX) units through a QC and normalization process developed and provided by Olink.

### Quality control for Olink plasma proteomics

The Olink PEA QC process consists of specifically engineered controls to monitor the performance of the main steps of the assays (immunoreaction, extension and amplification/detection) as well as the individual samples. Internal controls are spiked into each sample and represent a control using a non-human assay, an extension control composed of an antibody coupled to a unique DNA-pair always in proximity and, finally, a detection control based on a double stranded DNA amplicon. In addition, each plate run with Olink includes a control strip with sample controls used to estimate precision (intra- and inter-coefficient of variation). A negative control (buffer) run in triplicate is utilized to set background levels and calculate limit of detection (LOD), a plate control (plasma pool) is run in triplicate to adjust levels between plates, and a sample control (reference plasma) is included in duplicate to estimate CV between runs.

NPX is Olink’s relative protein quantification unit on a log2 scale and values are calculated from the number of matched counts on the NovaSeq run. Data generation of NPX consists of normalization to the extension control (known standard), log2-transformation, and level adjustment using the plate control (plasma sample).

### SomaScan plasma proteomic assays

The SomaScan Platform for proteomic profiling uses 4979 SOMAmer reagents, single-stranded DNA aptamers, to 4776 unique human protein targets. The modified aptamer binding reagents^32^, SomaScan assay^32,41^, its performance characteristics^29,42^, and specificity^43,44^ to human targets have been previously described. The assay used standard controls, including 12 hybridization normalization control sequences to control for variability in the Agilent readout process and 5 human calibrator control pooled replicates and 3 quality control pooled replicates to mitigate batch effects and verify the quality of the assay run using standard acceptance criteria.

### Quality control for SomaScan plasma proteomics

The SomaScan Assay is run using 96-well plates; 11 wells are allocated for control samples used to control for batch effects and to estimate the accuracy, precision, and buffer background of the assay over time. Five pooled Calibrator replicates, three pooled QC replicates, and three buffer replicates are run on every plate. The readout is performed using Agilent hybridization, scan, and feature extraction technology. Twelve Hybridization Control SOMAmers are added alongside SOMAmers to be measured from the biological samples and controls of each well during the SOMAmer elution step to control for readout variability. The control samples are run repeatedly during assay qualification and robust point estimates are generated and stored as references for each SOMAmer result for the Calibrator and QC samples. The results are used as references throughout the life of the SOMAscan V4 Assay. Plate Calibration is performed by calculating the ratio of the Calibrator Reference RFU value to the plate-specific Calibrator replicate median RFU value for each SOMAmer. The resulting ratio distribution is decomposed into a Plate Scale factor defined by the median of the distribution and a vector of SOMAmer-specific Calibration Scale Factors. Normalization of QC replicates and samples is performed using adaptive normalization by maximum likelihood (ANML) with point and variance estimates from a normal U.S. population. Post calibration accuracy is estimated using the ratio of the QC reference RFU value to the plate-specific QC replicate median RFU value for each SOMAmer. The resulting QC ratio distribution provides a robust estimate of accuracy for each SOMAmer on every plate. Plate-specific Acceptance Criteria: Plate Scale Factor between 0.4-2.5 and 85% of QC ratios between 0.8 and 1.2 must be met prior to release.

### Data analysis and visualization

All statistical analyses for the clinical and proteomics data in this cohort was performed using R version 4.0.2. All plots were generated using the ggplot2 package in R with the exception that the correlation plots were generated using the corrplot() function in R. Pairwise Pearson correlations were calculated for all proteins, and rows and columns of correlation plots were ordered based on hierarchical clustering. All heatmaps were generated using the heatmap3^45^ package and NPX values for each protein centered to have a mean of 0 and scaled to have a standard deviation of 1 within each protein. Scaled data greater than either 4 or 5 standard deviations from the mean were truncated at +/− 4 or 5. Rows and columns were ordered based on hierarchical clustering.

#### Unsupervised clustering

Principal components analysis (PCA) was performed using all proteins and all samples using the prcomp() function in R. Unsupervised clustering by UMAP was performed using all proteins, and either all samples or just day 0 samples, using the umap() function in R, and UMAP coordinates were plotted using the ggplot2 package. Unsupervised clustering by tSNE was by first performing dimensionality reduction by PCA and then taking the top principal components for a tSNE embedding using the Rtsne package and the argument pca=TRUE. k-nearest neighbor (KNN) graphs and Louvain community detection was performed using custom code and the FNN package provided in R.

#### Linear models

Linear regression models were fit independently to each protein using the lm package in R with protein values (NPX for Olink data) as the dependent variable. The models included a term for COVID-19 status and covariates for age, sex, ethnicity, heart disease, diabetes, hypertension, hyperlipidemia, pulmonary disease, kidney disease, immuno-compromised status to control for any potential confounding. P-values were adjusted to control the false discovery rate (FDR) at 5% using the Benjamini-Hochberg method implemented in the emmeans package in R.

#### Linear mixed models

Linear mixed effects models (LMMs) were fit independently to each protein using the lme4^46^ package in R with protein values (NPX for Olink data) as the dependent variable. The model for severity included a main effect of time, a main effect of severity, the interaction between these two terms, and a random effect of patient ID to account for the correlation between samples coming from the same patient. Covariates for age, sex, ethnicity, heart disease, diabetes, hypertension, hyperlipidemia, pulmonary disease, kidney disease, and immuno-compromised status were included in the model to control for any potential confounding. Significance of the three model terms was determined with an F-test using Satterthwaite degrees of freedom and type III sum of squares implemented with the lmerTest^47^ package in R. P-values for the three model terms of interest were adjusted to control the FDR at 5% using the Benjamini-Hochberg method. Group differences were calculated for each protein passing the FDR threshold with p-values adjusted using the Tukey method implemented by the emmeans package in R. Group differences with Tukey adjusted p-values less than 0.05 were considered statistically significant. Note, all other models were run similarly with time in addition to either Acuity_max_, age, or both age and severity as main effects instead of severity.

For SomaLogic data, LMMs for severity and time as main effects were run as was done for Olink. Overall, significant proteins were found to be partially overlapping with those found for Olink; for example, at D0, of the 1085 overlapping assays between the two platforms, 779 proteins were significant for severity or interaction term in Olink data, and 669 in the SomaLogic data, with 460 proteins overlapping between the two sets. The non-overlapping assays in part due to a narrower dynamic range for some of the SomaLogic assays.

#### Residuals

Model residual values were extracted from LMMs (as described above) independently fit to every protein using NPX as the dependent variable, age, sex, ethnicity, heart disease, diabetes, hypertension, hyperlipidemia, pulmonary disease, kidney disease, and immuno-compromised status as covariates and a random effect of patient ID to account for the correlation between samples taken from the same patient. These residuals represent the remaining unexplained variance in the protein expression after accounting for the effects of the included covariates.

#### Permutation controls

For the Olink assay, the likelihood of observing 1131 statistically significant proteins for the Acuity_max_ model term and 963 statistically significant proteins for the time and Acuity_max_ interaction term from the linear mixed models was evaluated using permutation testing. Acuity_max_ group was randomly permuted 100 times among patients and for each permutation the full LMM procedure was followed. None of the permutations produced as many statistically significant results as were observed when using the true Acuity_max_ groupings.

#### Gene set enrichment and pathway analysis

For analysis of functional pathways, two different strategies were employed: (i) gene set enrichment analysis^48^ using the ClusterProfiler package in R using the C7 immunologic signature gene set from the molecular signatures database v7.2 (https://www.gsea-msigdb.org/gsea/msigdb); and (ii) Ingenuity Pathway Analysis (Qiagen) on our gene lists using default parameters from the vendor. Pathways were visualized in dot plots and bar plots using the ggplot2 package in R.

#### Differential expression analysis of Louvain clusters

To identify plasma proteins that were differentially expressed between Louvain clusters, we created a linear model (as described above) with an identifier variable for Louvain group as a main effect and clinical variables (as described above) as covariates; that is, for proteins most significantly expressed in L1, our identifier variable was a 1 for L1 and 0 for all other groups. The top 25 differentially-expressed proteins were then visualized.

#### Prediction of severity

Predictive performance of severity within 28 days was performed using all proteins and model covariates and was estimated using elastic net logistic regression implemented by the glmnet^49^ package in R and 100 repeats of 5-fold cross validation. Model tuning was performed using the caret package in R. Variable scaling, model tuning, and feature selection was performed independently for each held-out fold such that the predictive model was never exposed to the held-out data. Measures of predictive performance are reported as medians and 95% confidence intervals calculated from the 100 repeats of the cross validation. Features were ranked by how frequently they were chosen to be included in the model.

#### Prediction of patient age and antibody neutralization level

Generalized linear models with lasso regularization were trained (using the R caret package) on COVID-19-positive patient proteome samples (consisting of 1472 Olink protein features) from each selected day (0, 3, and 7) to predict patient age (≤ or > 65 years) or neutralization levels (≤ or > 75%). For percent neutralization predictions, protein levels at day 0 were used to predict binned neutralization categories at day 3. Repeated 5-fold cross validation (with a hyperparameter scan from 0.0001 to 1 to select the lambda constant yielding the greatest prediction accuracy) was replicated 100 times to obtain a confidence interval for the area under the ROC curve (where ROC curves were generated using each patient’s estimated probability while serving as the held-out fold). The average feature weights of the final models from each of the 100 rounds of 5-fold cross validation were used to identify proteins of importance. Orthogonally, 10-fold cross validation was used to train and validate a random forest model (with default ntree=500 and mtry=38) to predict age (≤ or > 65 years) or neutralization quartiles (0-25, 25-50, 50-75, 75-100%) and important proteins were identified based on mean decrease in Gini. To identify protein features that were independent from or overlapping with severity markers, the union of the top 50 important features from the lasso and random forest models were intersected with significantly variable proteins between severity groups on day 0 (from the LMM described above).

### Measurement of neutralization levels

#### Constructs

SARS-CoV-2 S was amplified by PCR (Q5 High-Fidelity 2X Master Mix, New England Biolabs) from pUC57-nCoV-S (gift of Jonathan Abraham), in which the C-terminal 27 amino acids of SARS-CoV-2 S are replaced by the NRVRQGYS sequence of HIV-1, a strategy previously described for retroviruses pseudotyped with SARS-CoV S^50^. The truncated SARS-CoV-2 S fused to gp41 was cloned into pCMV by Gibson assembly to obtain pCMV-SARS2ΔC-gp41. psPAX2 and pCMV-VSV-G were previously described^51^. pTRIP-SFFV-EGFP-NLS was previously described^52^ (Addgene plasmid #86677). cDNA for human TMPRSS2 and the hygromycin resistance gene were generated by synthesis (Integrated DNA Technologies). pTRIP-SFFV-Hygro-2A-TMPRSS2 was generated by Gibson assembly.

#### Cell culture

293T cells were cultured in DMEM, 10% FBS (ThermoFisher Scientific), and PenStrep (ThermoFisher Scientific). 293T ACE2 cells (gift of Michael Farzan) were transduced with pTRIP-SFFV-Hygro-TMPRSS2 using TransIT®-293 Transfection Reagent (Mirus Bio, MIR 2700) to obtain 293T ACE2/TMPRSS2 cells, which were selected with 320 μg/ml of hygromycin (Invivogen) and used as a target in pseudotyped SARS-CoV-2 S lentivirus neutralization assays.

#### Pseudotyped SARS-CoV-2 lentivirus production and lentiviral production for transductions

The protocol for lentiviral production was previously described^51^. Briefly, 293T cells were seeded at 0.8 × 10^6^ cells per well in a 6-well plate and were transfected the same day with a mix of DNA containing 1 μg psPAX, 1.6 μg pTRIP-SFFV-EGFP-NLS, and 0.4 μg pCMV-SARS2ΔC-gp41 using TransIT®-293 Transfection Reagent. After overnight incubation, the medium was changed. SARS-CoV-2 S pseudotyped lentiviral particles were collected 30-34 hrs post medium exchange and filtered using a 0.45 μm syringe filter. To transduce 293T ACE2 cells, the same protocol was followed, with a mix containing 1 μg psPAX, 1.6 μg pTRIP-SFFV-Hygro-2A-TMPRSS2, and 0.4 μg pCMV-VSV-G.

#### SARS-CoV-2 S pseudotyped lentivirus antibody neutralization assay

The day before the experiment, 293T ACE2/TMPRSS2 cells were seeded at 5 × 10^3^ cells in 100 μl per well in 96-well plates. On the day of lentiviral harvest, 100 μl SARS-CoV-2 S pseudotyped lentivirus was incubated with 50 μl of plasma diluted in medium to a final concentration of 1:100. Medium was then removed from 293T ACE2/TMPRSS2 cells and replaced with 150 μl of the mix of plasma and pseudotyped lentivirus. Wells in the outermost rows of the 96-well plate were excluded from the assay. After overnight incubation, medium was changed to 100 μl of fresh medium. Cells were harvested 40-44 hrs post infection with TrypLE (Thermo Fisher), washed in medium, and fixed in FACS buffer containing 1% PFA (Electron Microscopy Sciences). Percentage GFP was quantified on a Cytoflex LX (Beckman Coulter), and data was analyzed with FlowJo.

### Processing and analysis of single-cell RNA-sequencing data

We analyzed 4 publicly available scRNAseq PBMC datasets from COVID-19 patients, which were obtained from: 1) Wilk *et al*, 2020^15^, COVID-19 atlas, https://www.covid19cellatlas.org/#wilk20; 2) Lee *et al*, 2020^8^, GEO accession GSE149689; 3) Arunachalam *et al*, 2020^4^, GEO accession GSE155673; and 4) Schulte-Schrepping *et al*, 2020^16^, EGA accession EGAS00001004571. Gene expression matrices after filtering low quality cells were used as provided by the respective investigators, and annotations were used as described in each of the studies. scRNAseq data from BAL fluid of COVID-19 patients from Bost *et al*, 2020^34^ was obtained from GEO accession GSE145926 and GSE149443. Cell-type specific expression in lung tissue was derived as described below. scRNAseq data from other tissues were obtained from the following sources: 1) heart from Tucker *et al*, 2020^53^, Broad Institute’s Single Cell Portal study ID SCP498; 2) kidney from Menon *et al*, 2020^54^, GEO accession GSE140989; 3) liver from MacParland *et al*, 2018^55^, GEO accession number GSE115469; and 4) pancreas from Baron *et al*, 2016^56^, expression matrix obtained from the Itai Yanai lab.

#### Generation of expression data from cell subsets in lung tissue

To generate lung cell-type specific signatures, we collected and aggregated scRNAseq studies, normalized each dataset, harmonized the published cell type annotations, and trained a multiclass logistic regression model.

##### Dataset selection

Only studies with scRNAseq data from primary tissue (including healthy, fibrotic, and COVID-19 donors), sequenced using the 10X Genomics platform, and published annotations were included. Two additional studies (chosen to maximize the number of cell types in the test set) were held out for cross validation to test cell type predictions and tune hyperparameters. The training datasets were Adams *et al*, 2020^57,58^, Chua *et al*, 2020^37^, Habermann *et al*, 2020^59^, Travaglini *et al*, 2020^60^, and two unpublished datasets. The test datasets used were Vieira *et al*, 2019^61^ and Laio *et al*, 2020^62^.

##### Normalization

Single cell/nuclei RNA-seq datasets from individual studies were aggregated and normalized using Scanpy^63^. Each study was subjected to identical pre-processing steps. First, UMI count values were winsorized, those above the 99th percentile of non-zero counts were reduced to the value of the 99th percentile (13 counts). Winsorized count data were normalized, so that UMI counts per cell/nucleus summed to 10,000, and then were logged, resulting in log(1+10,000*UMIs / total UMIs) for each cell/nuclei (“logtp10k”). Then the aggregated expression data were scaled using the scanpy ‘scale’ function with zero_center=False. To prepare cell type labels, we mapped each annotation to a common reference list before training. Cells labeled with cell types with ambiguous mappings (e.g. “T cell” or “myeloid”) were excluded from training.

##### Signature extraction

Cell type signatures were learned using an L2 penalized logistic regression model trained to predict the cell type from a single cell gene expression profile. The model was trained using SciKitLearn’s LogisticRegression function with the default parameters with the exception of C=.1, max_iter=30, and multi_class=‘ovr’. During fitting, individual cells were weighted to balance with respect to both cell type and study. Model coefficients learned were used as cell type signatures.

#### Analysis of scRNAseq data

All scRNAseq gene expression data was analyzed in R version 4.0.2 using custom code to look at average expression of genes of interest in each cell type. For visualization, gene expression was normalized across cell types (rows) with Z-scores and visualized in heatmaps using the heatmap3 function in R with hierarchical clustering of both cell types and genes. Where cell types were annotated on heatmaps, this was done by identifying cell types with the highest relative expression by Z-scores. The cell-type-specific intracellular gene list was defined as the top 20 genes with the highest relative expression for that cell type.

### Derivation of tissue-specific plasma proteomic signatures

Organ specific protein signatures were defined using RNA sequencing data from the Genotype-Tissue Expression (GTEx) Portal (https://www.gtexportal.org/home/). The median transcripts per million (TPM) of 56,200 genes across 54 non-diseased tissue sites were obtained. For each tissue site, the intersection of the top 500 highest TPM genes and the top 500 most variable genes (based on coefficient of variation across tissue types) was identified (**Table S14**). Proteins that were also measured by SomaScan were extracted, validated for high tissue specific expression, and consolidated across related tissues for each organ of interest. Organ signatures were split based on localization (intracellular versus membrane/secreted) using UniProt and literature annotations. The values for each protein across all COVID-19-positive patients were scaled to Z scores, and the mean Z score of all proteins in an organ set was used as an overall signature score for a given patient.

### Ligand-receptor analysis

Single-cell RNAseq expression profiles (10X genomics) of immune cells isolated from BAL fluid of healthy and COVID-19-infected patients of varying severity from Bost *et al*, 2020^34^ was obtained from GEO accession GSE145926 and GSE149443. Python 3.8 was used to run the python package Cellphonedb v2.1.4 with the following parameters: database v2.0.0, statistical method analysis, 1000 iterations, 6000 cell subsampling. The metadata cluster identities were previously assigned based on the published annotations. Analysis of specific ligands and receptors was performed from a curated list of known ligand-receptor pairs, and cell types were assigned to particular ligands and receptors by identifying cell types with the highest relative expression by Z-scores.

## Data and code availability

Data for the Olink proteomics assay (1352 proteins screened in 384 patients) is publicly available at https://www.olink.com/mgh-covid-study/. All other data and code will be made available at the time of publication.

## Extended Data Figure Legends

**Extended Data Figure 1.**
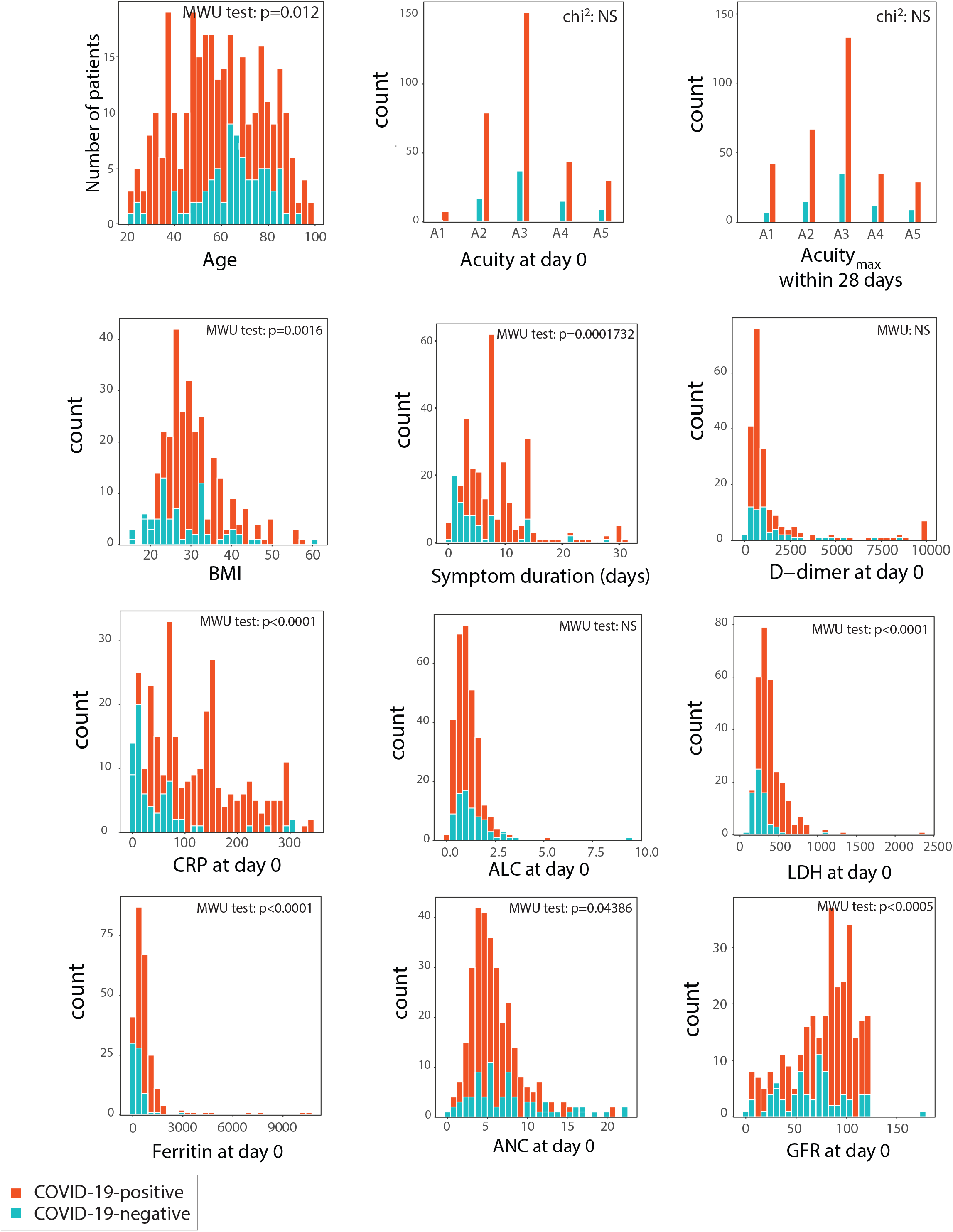
Distributions of clinical characteristics of the study cohort. Age, disease acuity, Acuity_max_, BMI, symptom duration prior to ED arrival, D-dimer, CRP, absolute lymphocyte count (ALC), LDH, ferritin, absolute neutrophil count (ANC), and glomerular filtration rate (GFR). All measurements, except Acuity_max_, symptom duration, and BMI, are at day 0. Significance is by the Mann-Whitney U-test (MWU) and chi-squared test (chi^2^); p-values as indicated.

**Extended Data Figure 2.**
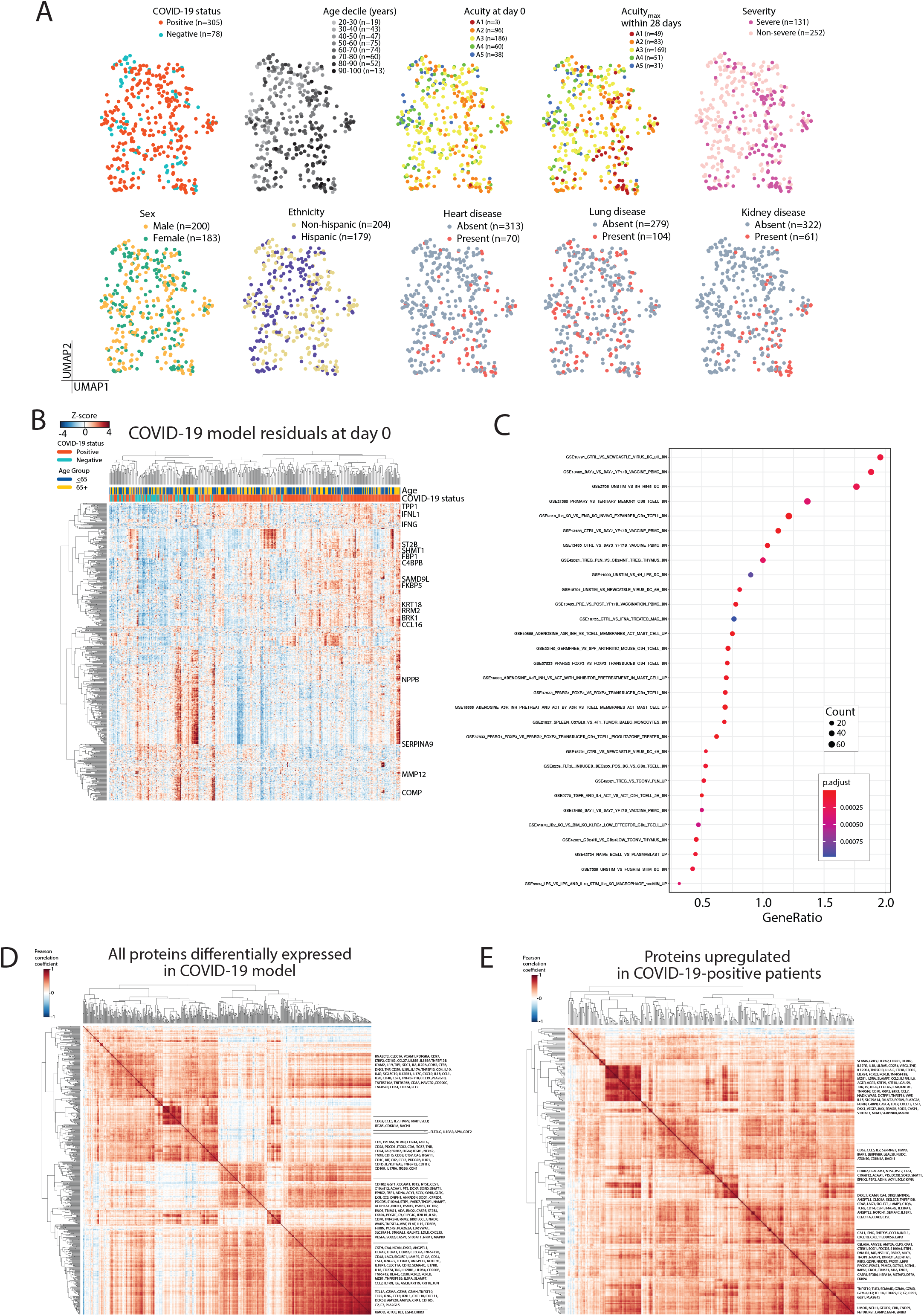
Differentially-expressed plasma proteins between COVID-19-positive and negative patients. (A) Unsupervised clustering (by UMAP), generated from plasma proteins of all patients on day 0, color-coded (left to right) by COVID-19 status, age decile, acuity level at day 0, Acuity_max_ within 28 days, severity, sex, ethnicity, previously known heart disease, previously known lung disease, and previously known kidney disease. (B) Identification of differentially-expressed proteins by COVID-19 status using linear model of Olink proteins with putative confounders as covariates (see **online methods**). Heatmap of residuals from this model for each protein found to be significantly differentially-expressed between COVID-19-positive and negative patients, using the model described in **Fig. 1D** and in (C)-(E) below. (C)-(E) Differentially-expressed proteins by COVID-19 status. Linear model fitting each Olink protein, with COVID-19 status as a main effect and putative confounders as covariates (see **online methods**). p-values were adjusted to control the false discovery rate (FDR) at < 0.05, Benjamini-Hochberg method. (C) Gene-set enrichment analysis for pathways enriched among plasma proteins differentially-expressed between COVID-19-positive and negative patients. (D) Correlation heatmap of plasma protein levels of all proteins significantly differentially-expressed between COVID-19-positive and negative patients. (E) Correlation heatmap of plasma protein levels of all proteins significantly higher in COVID-19-positive patients than in COVID-19-negative patients.

**Extended Data Figure 3.**
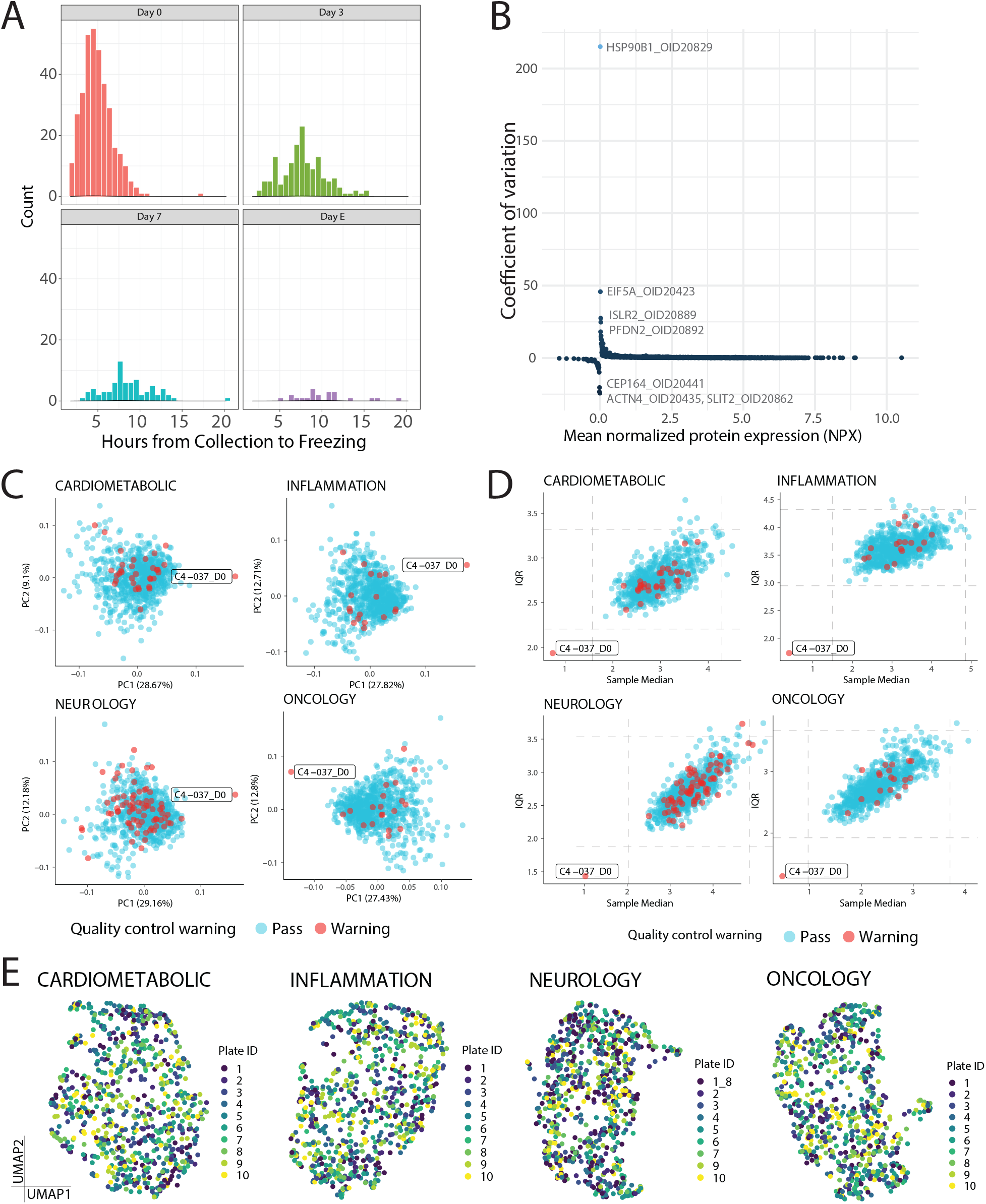
Quality control for Olink plasma proteomics data. (A) Histogram showing the time from collection to freezing for each sample, separated by time point of collection. E, event-driven samples (see **online methods**). (B) Histogram showing the coefficient of variation versus mean normalized protein expression (NPX) for each of the 1356 assays performed by Olink. (C) Principal component plots using Olink assays in four predefined Olink panels (cardiometabolic, inflammation, neurology, and oncology), each with >300 protein assays. Shown are first two principal components (PC1 vs. PC2) for all patient samples. (D) Scatterplot showing interquartile range (IQR) versus sample median for each sample. In (C) and (D), sample C4-037_D0 was noted to be an outlier and was therefore excluded from downstream analyses. (E) Unsupervised clustering by UMAP of all samples by assays in each predefined panel as in (C-D), color-coded by plate label, confirming there were no plate-related biases.

**Extended Data Figure 4.**
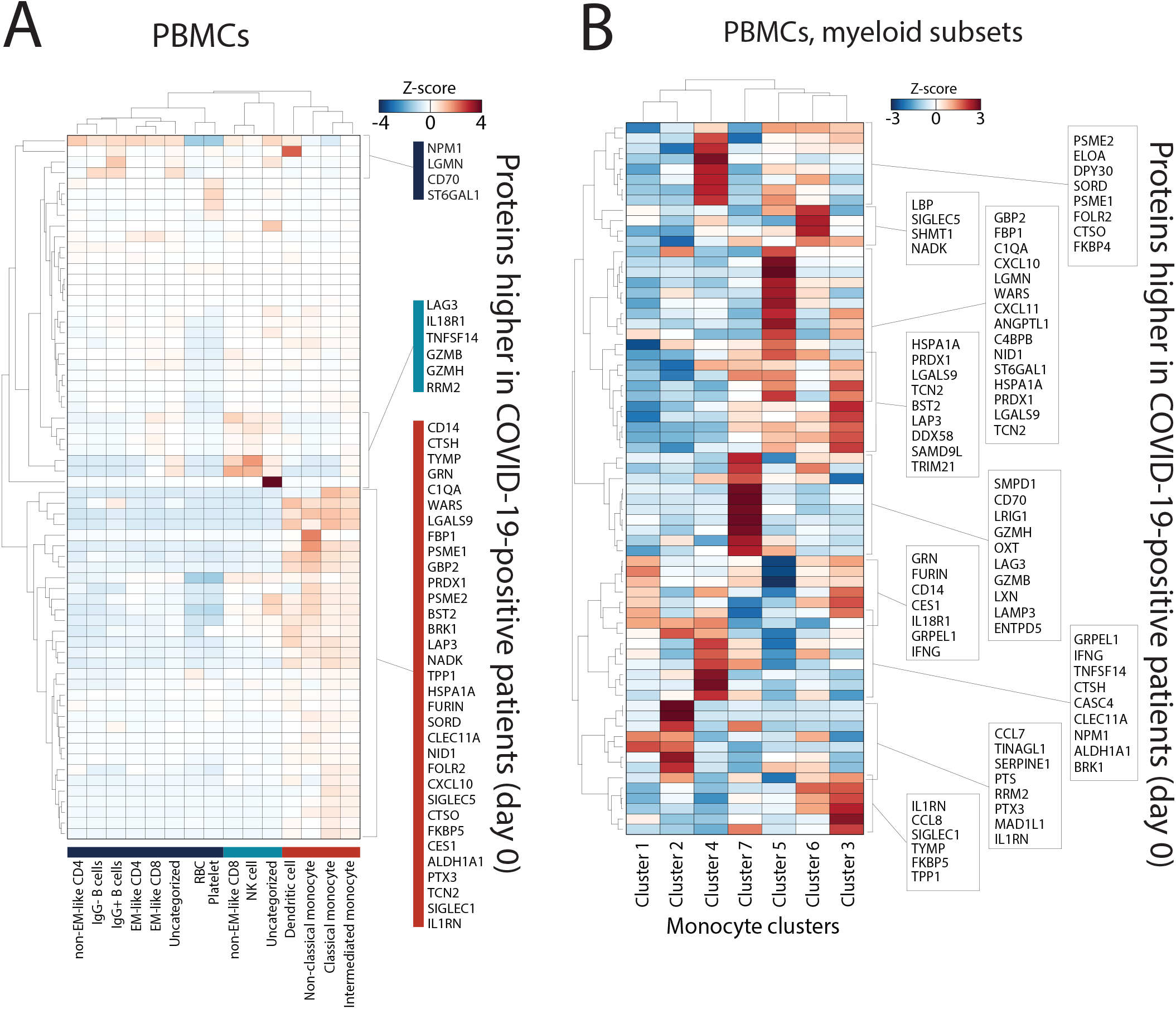
Inference of cell of origin by mapping gene expression of differentially-expressed plasma proteins (elevated in COVID-19-positive versus COVID-19-negative patients) onto scRNAseq peripheral blood cell COVID-19 datasets. Heatmaps of mean expression of COVID-19-related proteins (y-axis) in immune cell subtypes (x-axis) from (A) Lee *et al*.^8^ and (B) Schulte *et al*.^16^

**Extended Data Figure 5.**
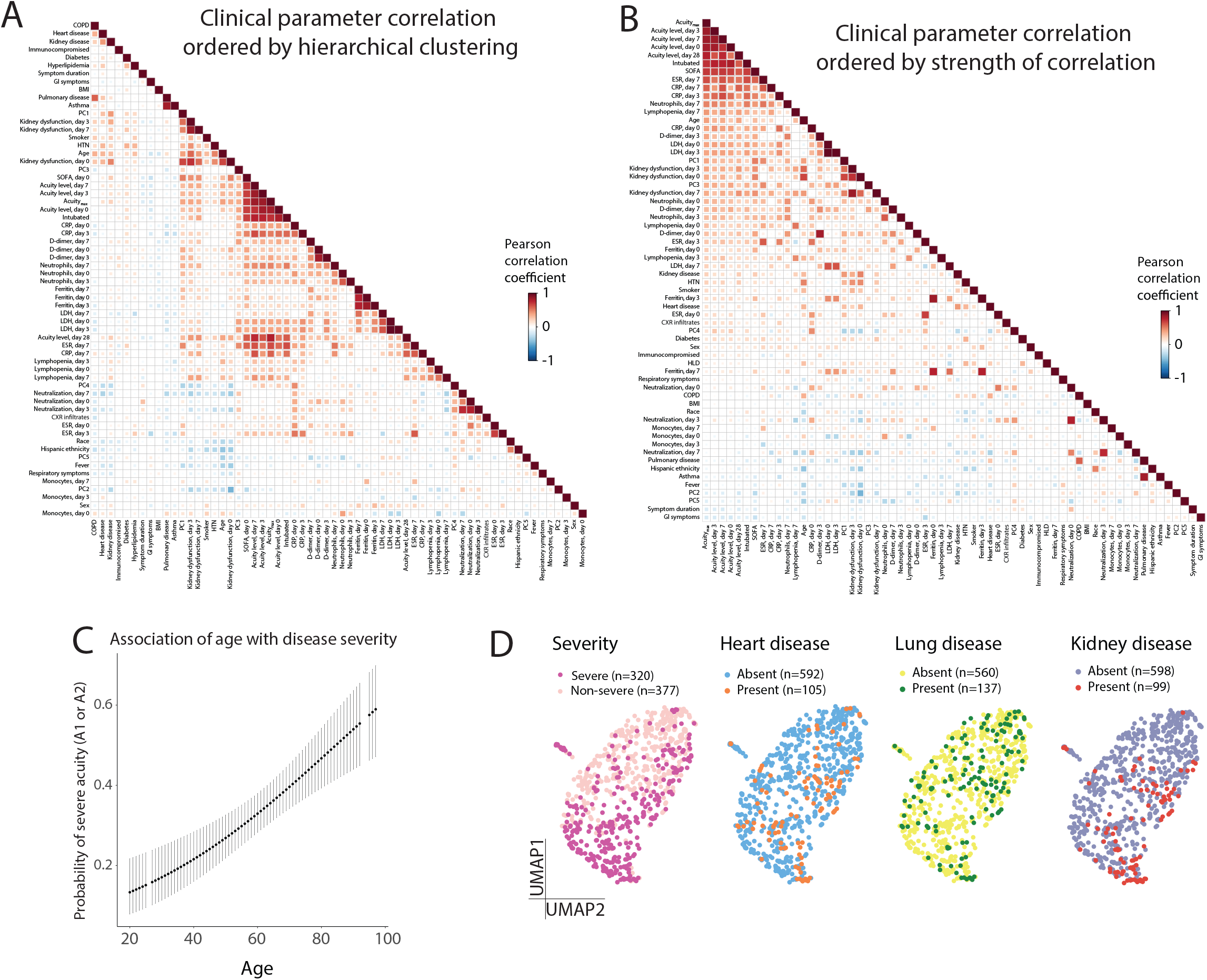
Clinical correlates for patients in this study cohort. (A)-(B) Correlation heatmap of all clinically associated variables collected in this study ordered by hierarchical clustering (A) or correlation to Acuity_max_ (B). For the purpose of displaying positive correlation with increasing disease severity, inverse numbering was assigned to A1-A5 for these calculations, and kidney function was measured as the inverse of the glomerular filtration rate (GFR). PC1 to PC5 are the loadings of the first five principal components for the PCA plot of all samples using all proteins. (C) Logistic regression model predicting probability of severe acuity (Acuity_max_ of A1 or A2) using patient age. (D) Unsupervised clustering by UMAP using plasma proteins of all COVID-19-positive patients at days 0, 3 and 7, color-coded (left to right) by severity, previously known heart disease, previously known lung disease, and previously known kidney disease.

**Extended Data Figure 6.**
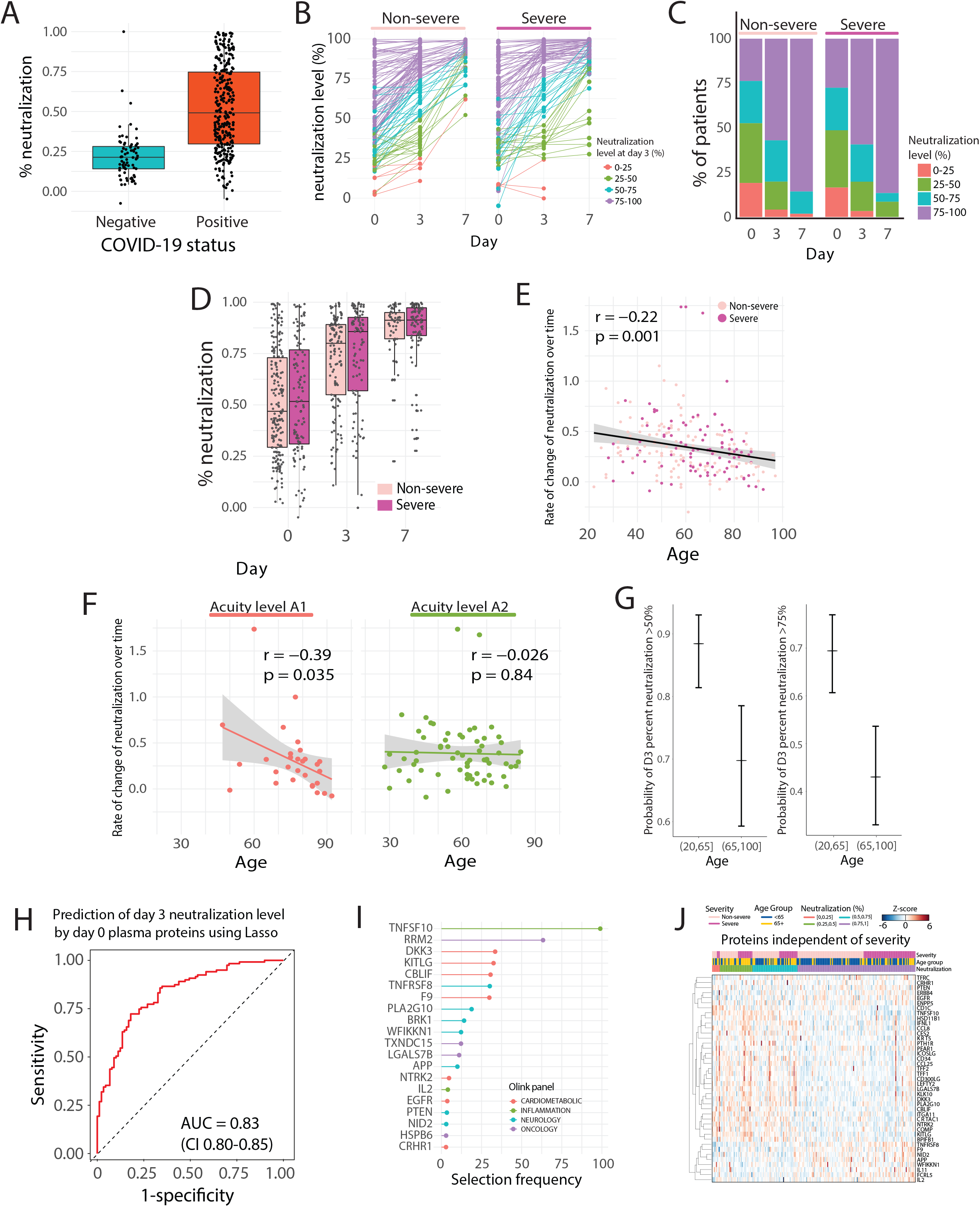
Predictors of neutralization and its association with disease severity and age. (A) Box plot showing neutralization levels for COVID-19-negative and positive patients at day 0. Box edges mark the interquartile range (IQR) and the line in the middle is the median. (B) Point-range plots showing neutralization levels in non-severe and severe COVID-19-positive patients over time. Color coding is by neutralization level at day 3, grouped into 0-25%, 25-50%, 50-75%, and 75-100%. (C) Proportion of non-severe and severe patients within each of the day 3 bins defined in (B) over time. (D) Box plots showing neutralization levels in non-severe and severe patients over time. Box edges mark the IQR and the line in the middle is the median. (E) Scatter plot showing the correlation of age with the rate of change in neutralization level in COVID-19-positive patients over time from day 0 to day 7. Rate of change is the negative of the regression line slope through log2(fold change) in GFP levels at each timepoint compared to controls. (F) Scatter plot showing the correlation of age with rate of change in neutralization level over time in A1 patients (left) and A2 patients (right). (G) Logistic regression model predicting neutralization titers above 50% (left) or above 75% (right), stratified by young (≤65 yrs) and elderly (>65 yrs) age. (H) Lasso regression model for prediction of day 3 neutralization level (above or below 75%) using Olink plasma proteins at day 0 across all COVID-19-positive patients. The prediction was performed with 5-fold cross-validation over 100 iterations; AUC 0.83 (95% CI 0.80-0.85). (I) Selection frequency of the top selected features for the neutralization predictor in (H). (J) Heatmap showing the Olink plasma protein expression of each of the top selected features from the predictor in (H) that did not overlap with the top severity-associated proteins from the linear mixed model described in **Fig. 2D**.

**Extended Data Figure 7.**
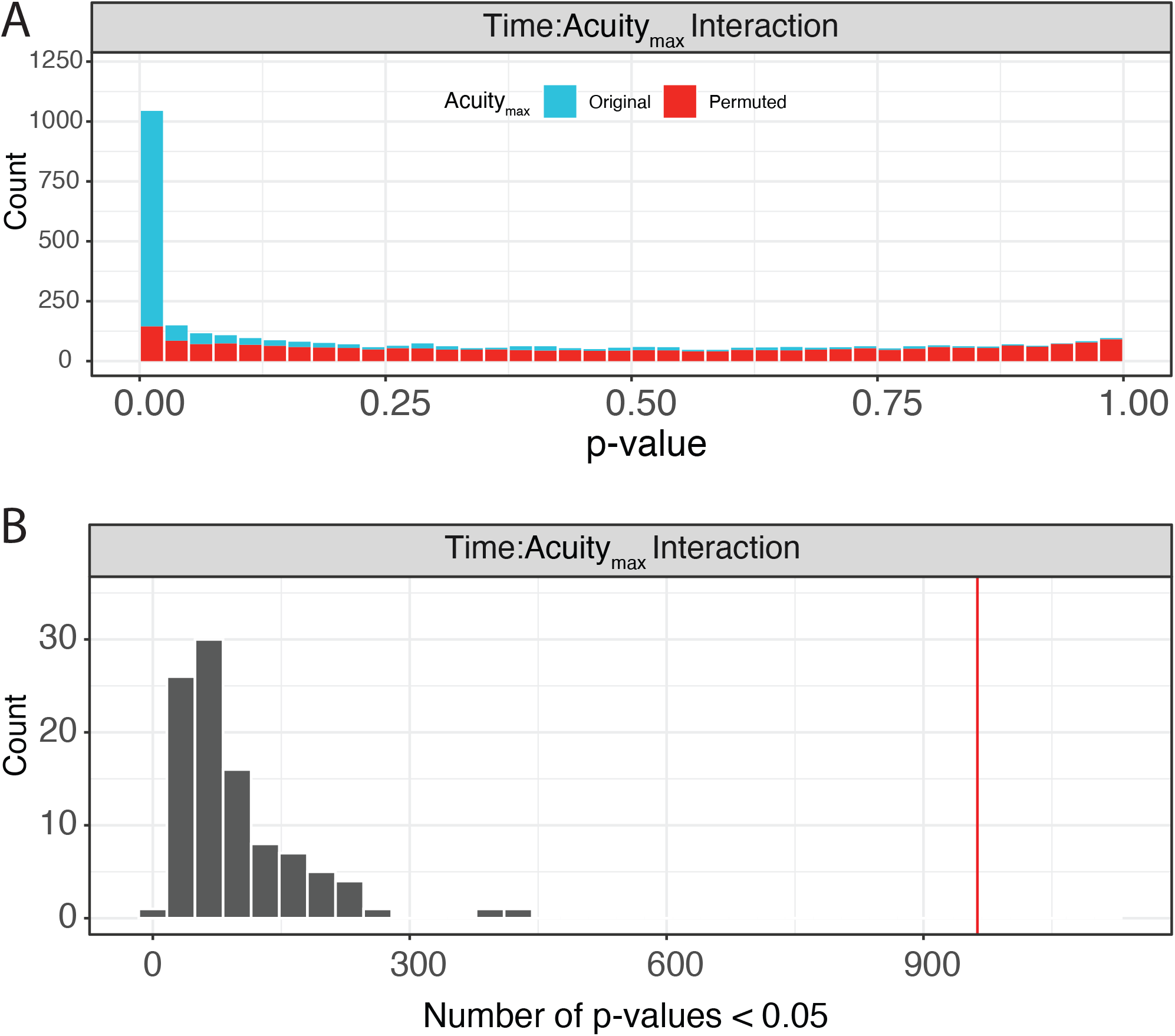
Permutation analysis for linear mixed model on Olink data, with Acuity_max_ and time as main effects. (A) Linear mixed model fitting each Olink protein, with Acuity_max_, timepoint, and the interaction of the two terms as main effects and putative confounders as covariates (see **online methods**). Significance of the three model terms was determined with an F-test using Satterthwaite degrees of freedom and type III sum of squares. p-values for the three model terms of interest were adjusted to control the FDR at < 0.05, Benjamini-Hochberg method. Group differences were calculated for each protein passing the 0.05 FDR threshold with p-values adjusted using the Tukey method. Shown are the distribution of p-values obtained from this linear mixed model and the distribution of p-values obtained from permutations of random Acuity_max_ assignments to all patients. This result shows that the obtained distribution of p-values does not happen by chance. (B) Distribution of the number of p-values < 0.05 obtained from 100 permutations of random Acuity_max_ assignments for the linear mixed model described in (A). The red line indicates the number of significant assays obtained from our true model with correct patient Acuity_max_ assignments.

**Extended Data Figure 8.**
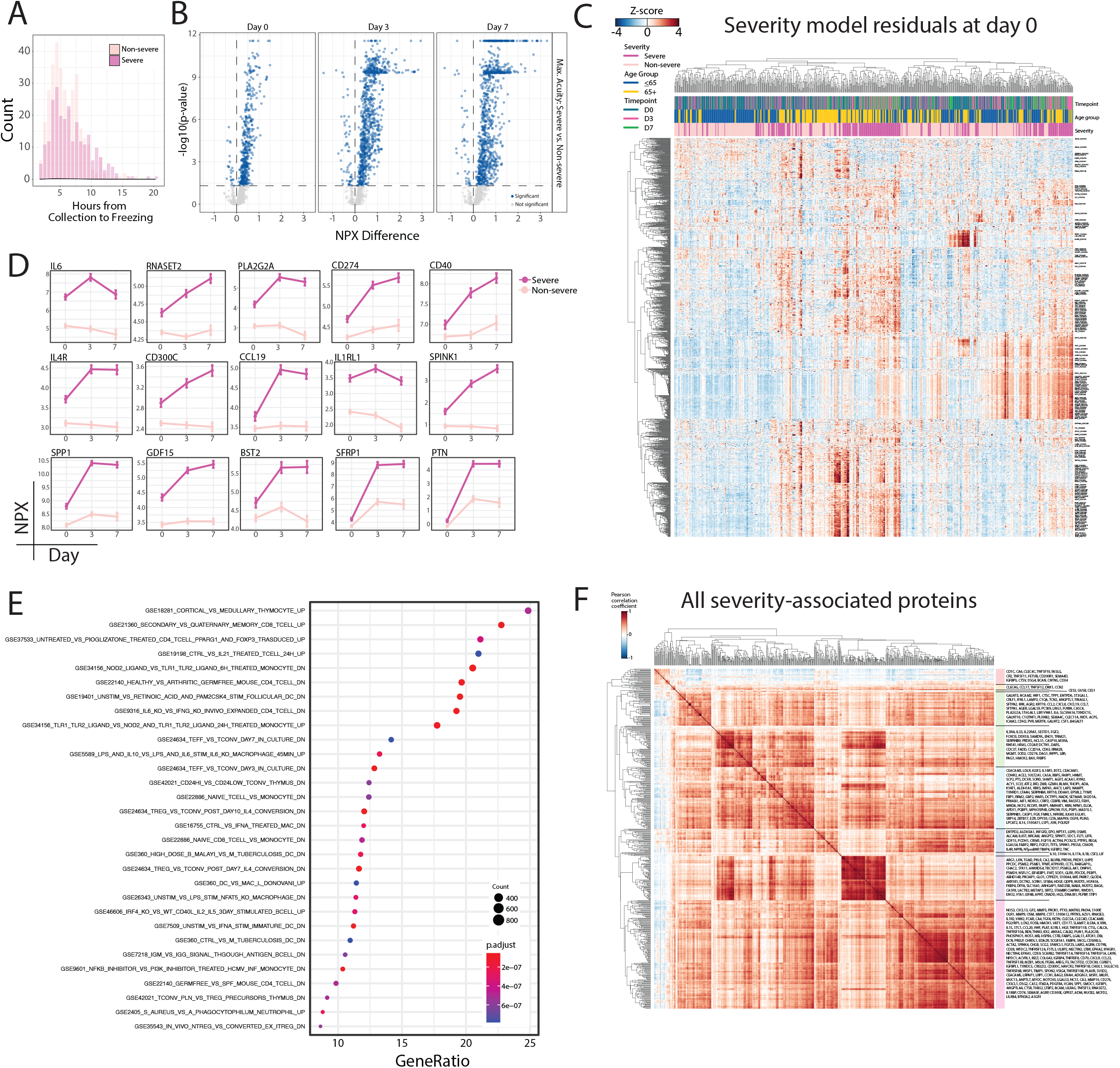
Protein associations with disease severity. (A) Time from sample collection to freezing for all samples, color-coded by COVID-19 severity, indicating no clear difference between samples from severe and samples from non-severe patients. (B) Linear mixed model fitting each Olink protein, with severity, timepoint, and the interaction of the two terms as main effects and putative confounders as covariates, as in **Fig. 2D** (see **online methods**). Volcano plots of differentially-expressed proteins between severe and non-severe COVID-19-positive patients at each time point. Blue circles, proteins that are significantly differentially-expressed. (C) Linear model fitting each Olink protein, with putative confounders as covariates (see **online methods**). Heatmap of residuals from this model for each protein that was found to be significant for interaction term in the model described in **Fig. 2E**. (D) Point range plots over time of a selected set of proteins significant for interaction term in the model described in (B), color-coded by disease severity. (E) Gene-set enrichment analysis for pathways enriched among plasma proteins differentially-expressed between severe and non-severe COVID-19-positive patients. (F) Correlation heatmap of plasma protein expression levels of all proteins significantly differentially-expressed between severe and non-severe COVID-19-positive patients.

**Extended Data Figure 9.**
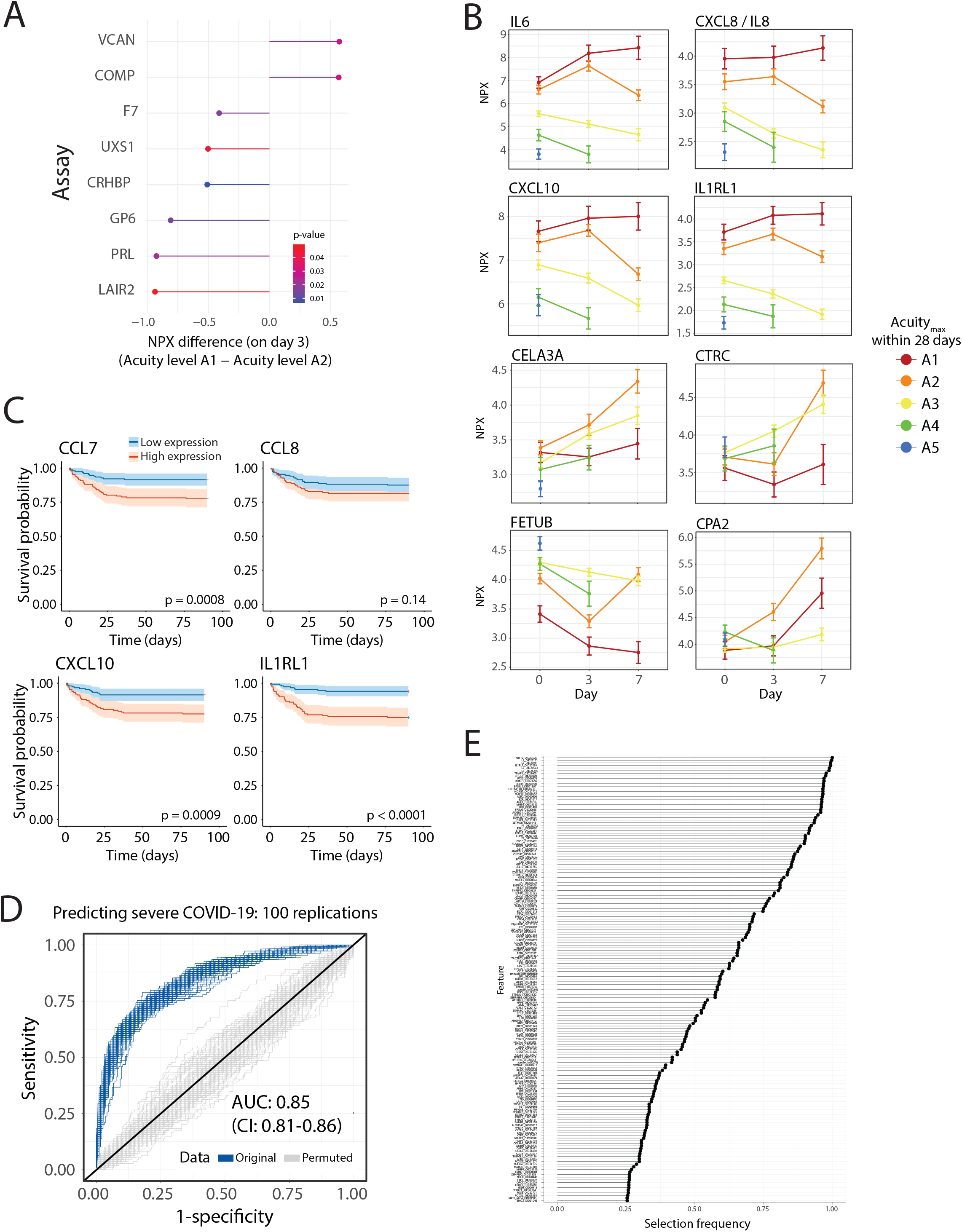
Protein associations with death in ARDS patients. (A) Differentially-expressed proteins at day 3 between patients who had a maximum classification of A1 (death) versus A2 (ARDS but survived) within day 28. Derived from linear mixed model fitting each Olink protein, with Acuity_max_ within day 28, timepoint, and the interaction between the two terms as main effects, as in **Fig. 2E**. (B) Point-range plots for select proteins from the model in (A). (C) Kaplan-Meier curves for overall survival of patients stratified by higher or lower than median expression of CCL7, CCL8, CXCL10, or IL1RL1. (D) Receiver operating characteristic (ROC) curve showing predictive performance of an elastic net logistic regression classifier of disease severity using Olink plasma proteins for each patient at day 0. Shown are curves for each of the 100 repeats of 5-fold cross validation. Measures of performance are reported as medians and 95% confidence intervals. Shown in the figure are the ROC curves for the original data with the true severity labels and the curves for our data with random (permuted) severity assignments. (E) Frequency of protein feature selection over the 100 iterations of the predictor in (D).

**Extended Data Figure 10.**
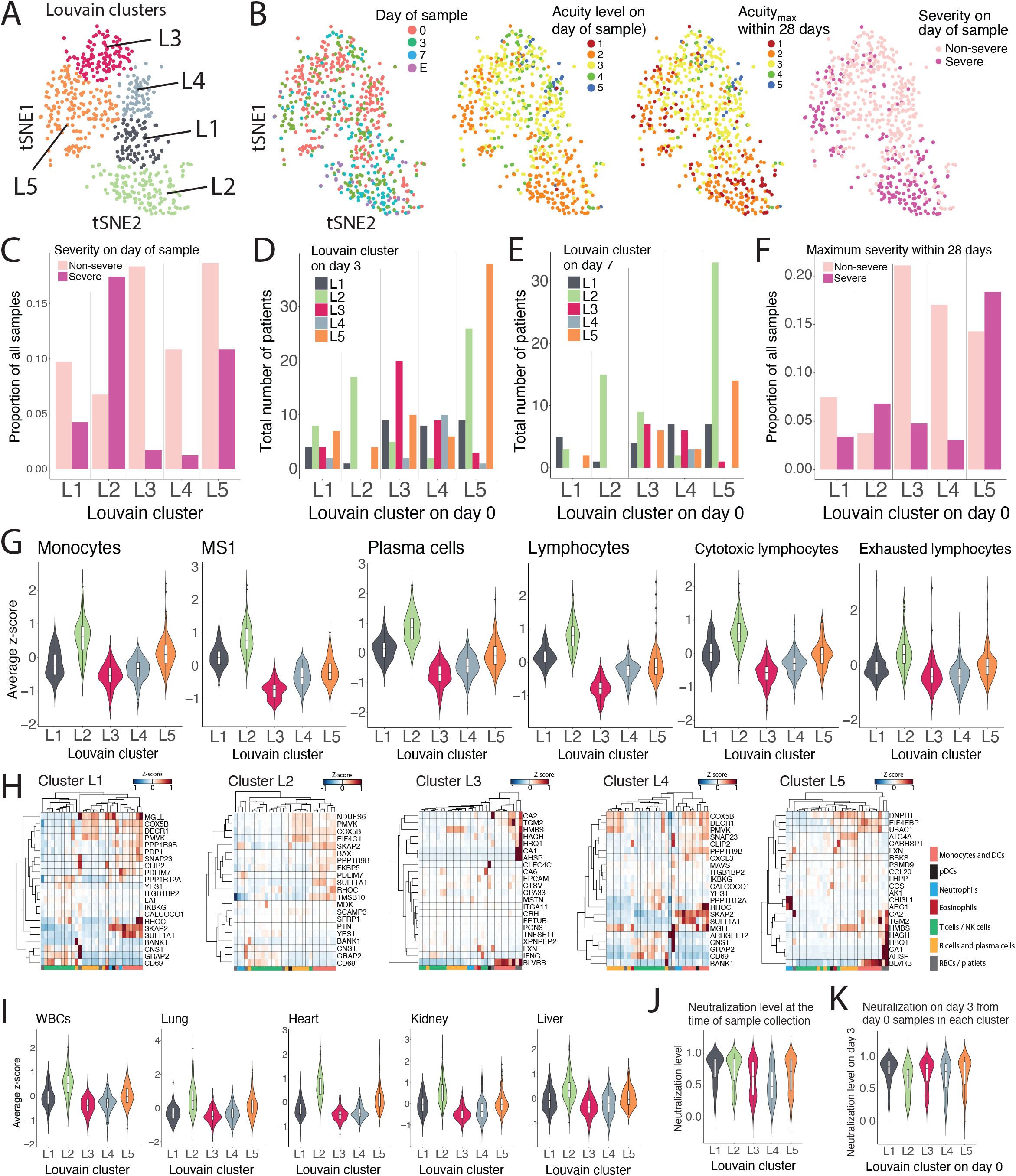
Unsupervised patient clustering using Olink plasma proteomics data. (A) Unsupervised clustering (using t-statistic stochastic network embedding, tSNE) of all patient samples using all Olink plasma proteins with graphical clustering using the Louvain algorithm. The optimal solution was five distinct clusters, labeled L1 to L5. (B) tSNE plots as in (A) labeled (left to right) by day of sample collection, acuity level at day of sample, Acuity_max_ within day 28, and disease severity. E, event-driven samples (see **online methods**). (C) Distribution of severity on the day of sample collection for samples in each Louvain cluster in (A). (D) Louvain cluster on day 3 for the day 0 samples from each of the Louvain clusters. (E) Louvain cluster on day 7 for the day 0 samples from each of the Louvain clusters. (F) Maximum severity within day 28 in each of the Louvain clusters in (A). (G) Violin plots showing expression within each Louvain cluster of cell-type specific plasma proteins, derived from gene expression data, for monocytes, MS1 monocytes, plasma cells, lymphocytes, cytotoxic lymphocytes, and exhausted lymphocytes. (H) Differentially-expressed proteins in each cluster compared to all other patient samples. Shown next to each protein is the gene expression within scRNAseq of PBMCs of COVID-19-positive patients^15^. Cell types are color-coded. (I) Violin plots showing expression of tissue-specific signatures, derived as in **Fig. 3A**, in each Louvain cluster. (J) Violin plot showing neutralization level at the time of sample collection for each Louvain cluster. (K) Violin plots showing neutralization at day 3 for all day 0 samples in each Louvain cluster.

**Extended Data Figure 11.**
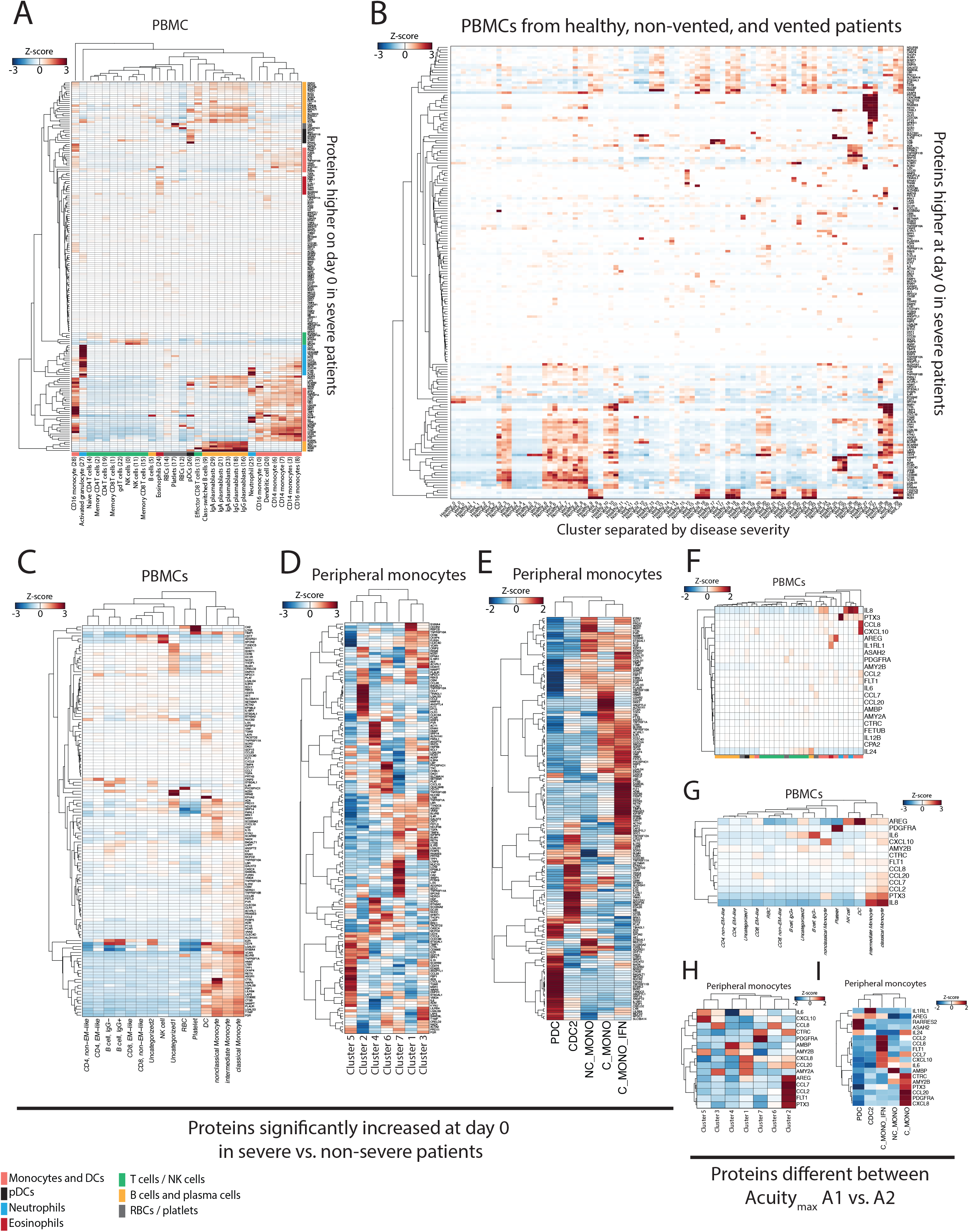
Expression of severity-associated plasma proteins in PBMCs from COVID-19 patients. (A) Gene expression of Olink plasma proteins expressed more highly in severe COVID-19 patients shown within PBMCs using scRNAseq data^15^. (B) Protein expression as in (A), with cell types separated by healthy, and COVID-19-positive patients, segregated by disease severity. (C)-(E) Expression of Olink plasma proteins expressed more highly in severe COVID-19 patients within PBMCs show within another independent dataset^8^ (C), or circulating monocytes, from two independent datasets^4,16^ (D)-(E). (F)-(I) Expression of differentially-expressed Olink proteins at day 7 between patients whose maximum classification was A1 (death) versus A2 (ARDS but survived) within day 28, for two independent PBMC datasets^8,15^ ((F)-(G)) and two independent peripheral monocyte datasets^4,16^ ((H)-(I)).

**Extended Data Figure 12.**
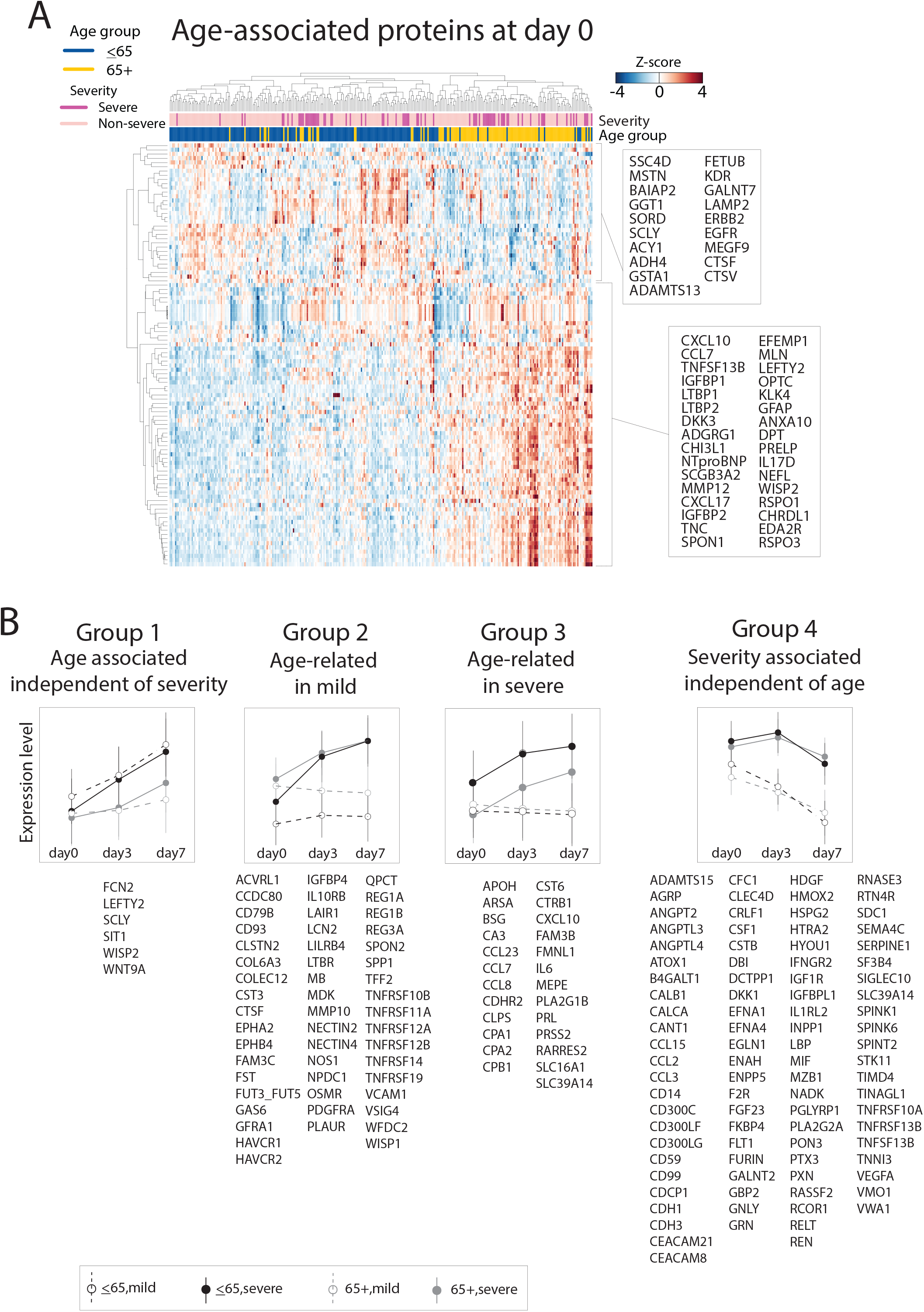
Proteins implicated in severity and aging. (A) Linear mixed model with age group (≤65 versus >65 years) and time point as main effects and all other clinical variables as covariates, as above, for Olink proteins. Heatmap of top 100 differentially-expressed proteins at D0. (B) Linear mixed model fitting each Olink protein, with age group, time point, disease severity and interaction terms as main effects and covariates, as in panel. Examples are shown of distinct patterns of expression seen in aged and young patients, with non-severe (mild) and severe disease.

**Extended Data Figure 13.**
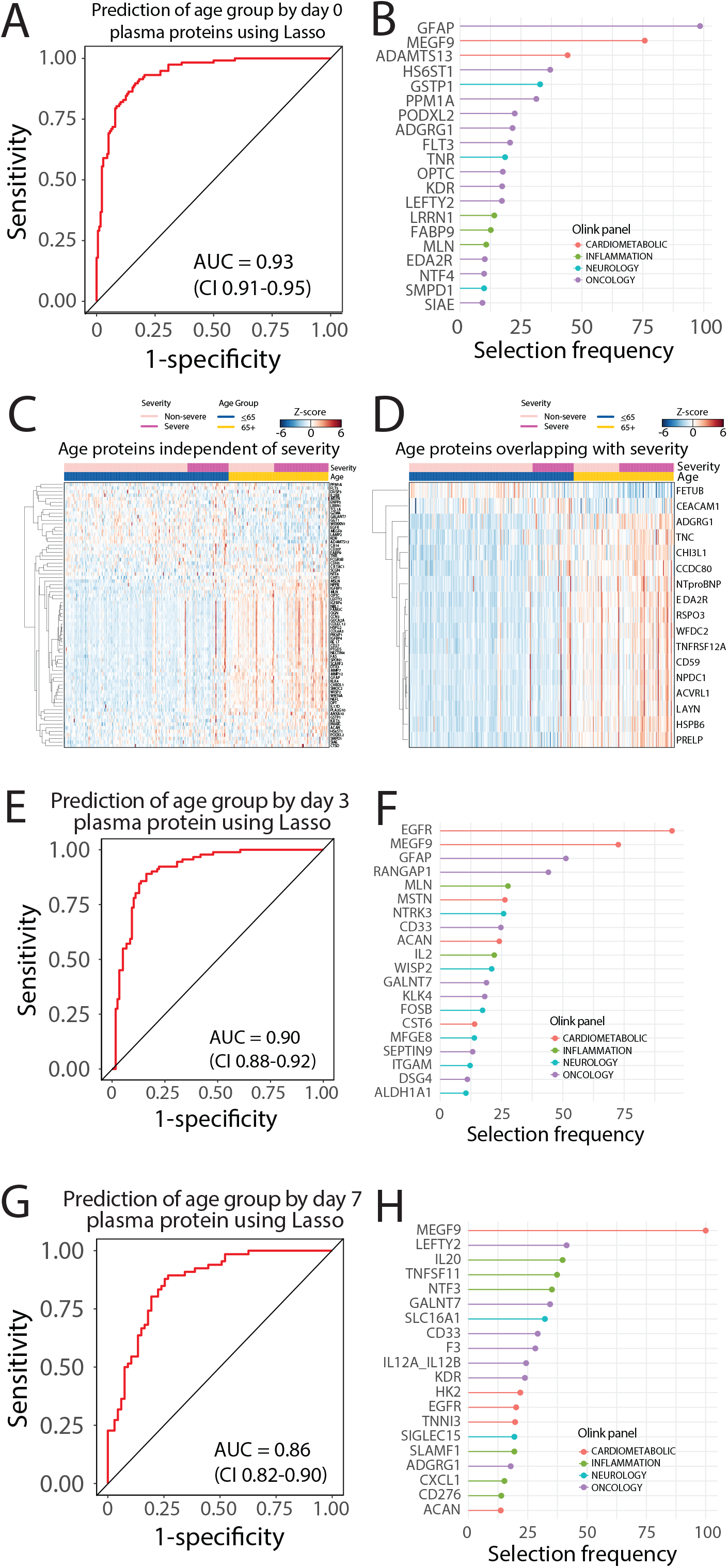
Prediction of patient age. (A) Lasso regression model for prediction of patient age group using Olink plasma proteins at day 0 across all COVID-19-positive patients. Performed with 5-fold cross-validation over 100 iterations; AUC 0.93 (95% CI 0.91-0.95). (B) Selection frequency of the top selected features for age predictor in (A). (C)-(D) Plasma protein expression heatmaps of each of the top selected features from the predictor in (A) that did not overlap (C) or did overlap (D) with the top severity-associated proteins from the linear mixed model described in **Fig. 2D**. (E) Lasso regression model for prediction of patient age group using Olink plasma proteins at day 3 across all COVID-positive patients. Performed with 5-fold cross-validation over 100 iterations; AUC 0.90 (95% CI 0.88-0.92). (F) Selection frequency of the top selected features for the age predictor in (E). (G) Lasso regression model for prediction of patient age level using Olink plasma proteins at day 7 across all COVID-positive patients. Performed with 5-fold cross-validation over 100 iterations; AUC 0.86 (95% CI 0.82-0.90). (H) Selection frequency of the top selected features for the age predictor in (G).

**Extended Data Figure 14.**
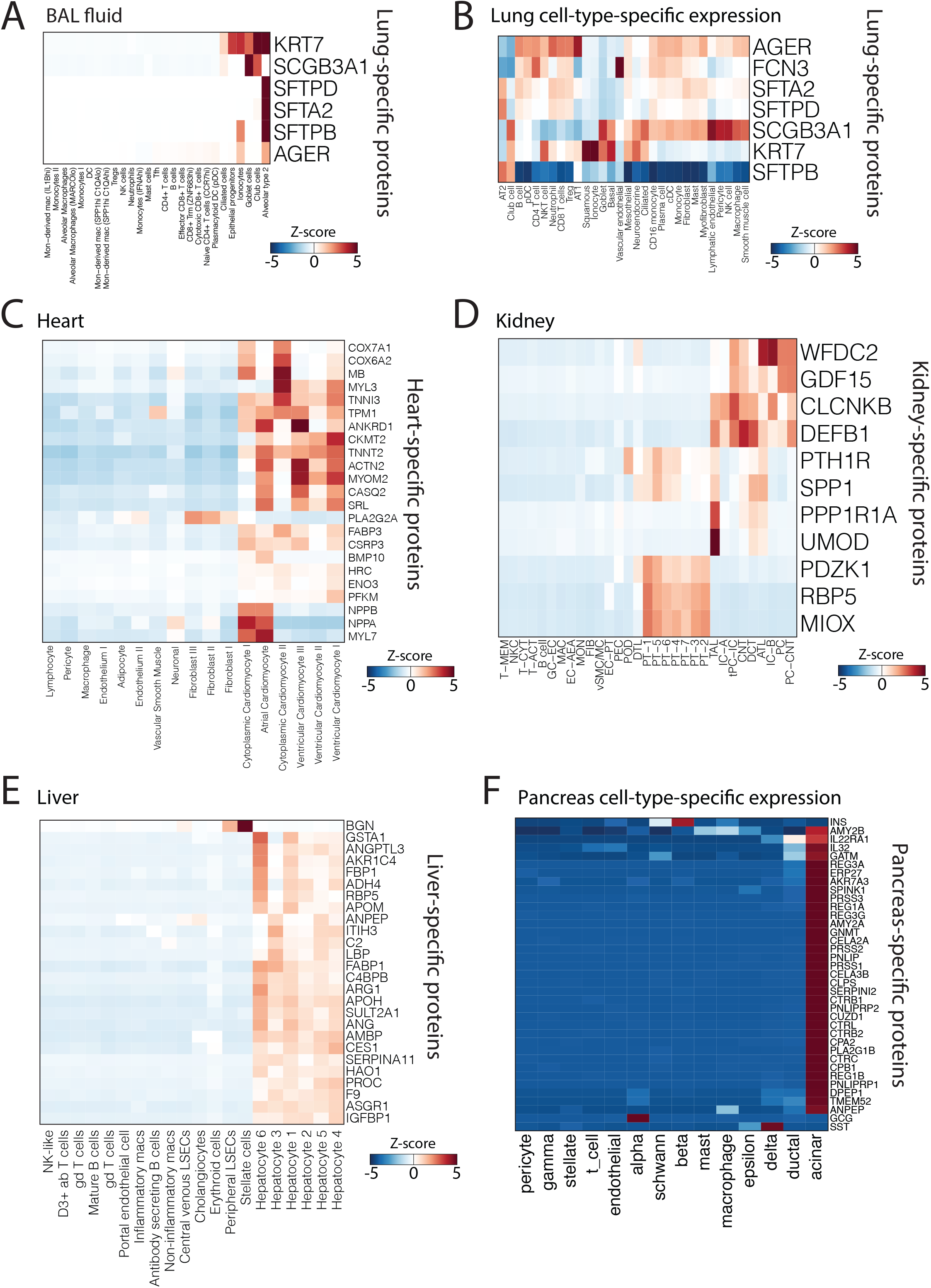
Tissue-specific cell-type expression of organ damage plasma protein signatures. (A)-(B) Expression of derived lung specific plasma proteins within (A) BAL fluid^34^ from COVID-19 patients and (B) an integrated analysis of several lung scRNAseq datasets. (C) Expression of derived heart specific plasma proteins within specific cell subsets from cardiac single-nucleus RNA-seq data^53^. (D) Expression of derived kidney specific plasma proteins within specific cell subsets from kidney scRNAseq data^54^. (E) Expression of derived liver specific plasma proteins within specific cell subsets from liver scRNAseq data^55^. (F) Expression of derived pancreas specific plasma proteins within specific cell subsets from a pancreatic scRNAseq dataset^56^.

**Extended Data Figure 15.**
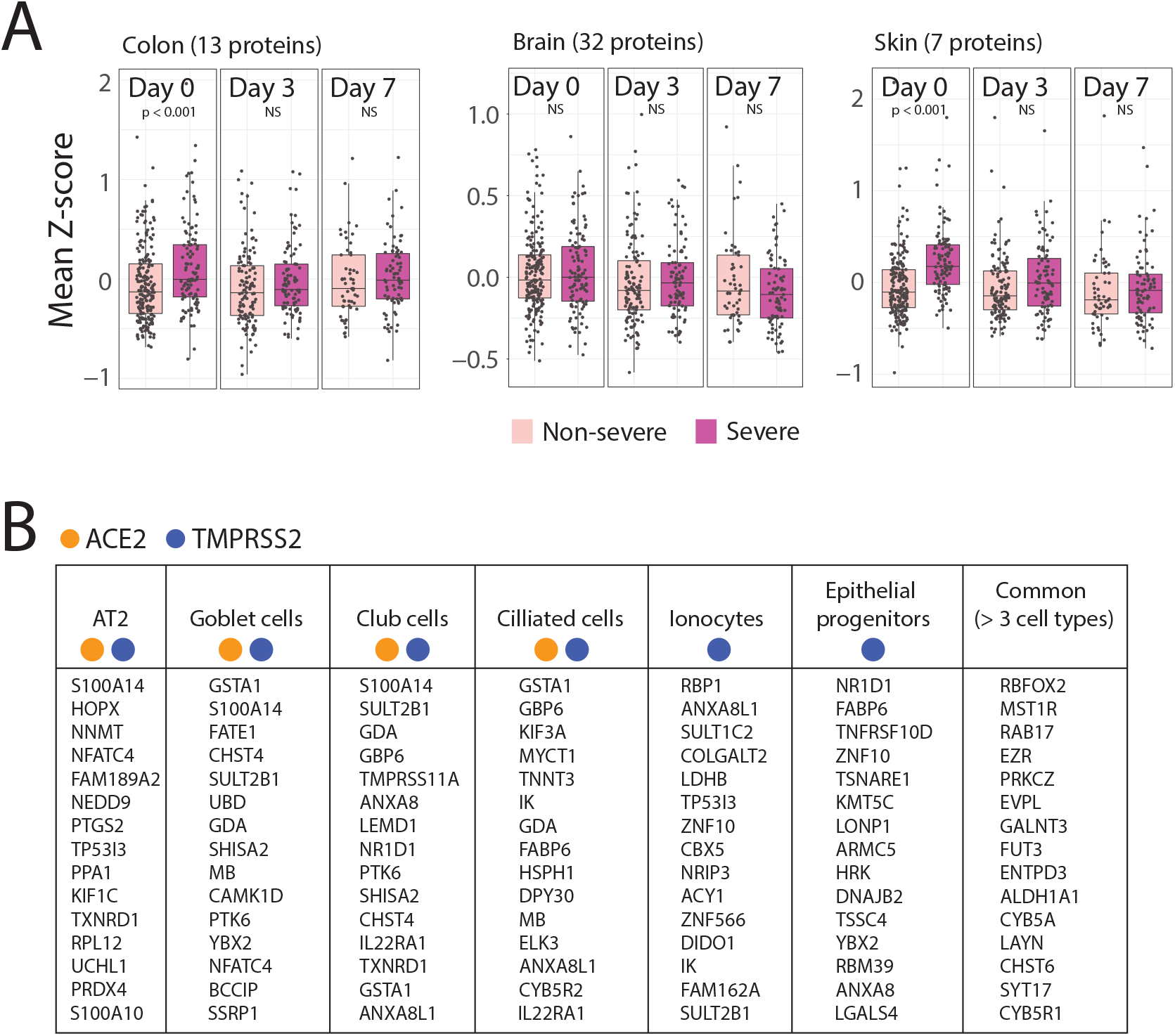
Organ-specific cell death signatures. (A) Expression of tissue-specific plasma protein signatures in severe versus non-severe patients at each timepoint in select tissues. (B) Top differentially-expressed genes per cell type obtained from the subset of severity-associated intracellular plasma proteins at D0. ACE2 and TMPRSS2 expression indicated by orange and blue circles, respectively.

**Extended Data Figure 16.**
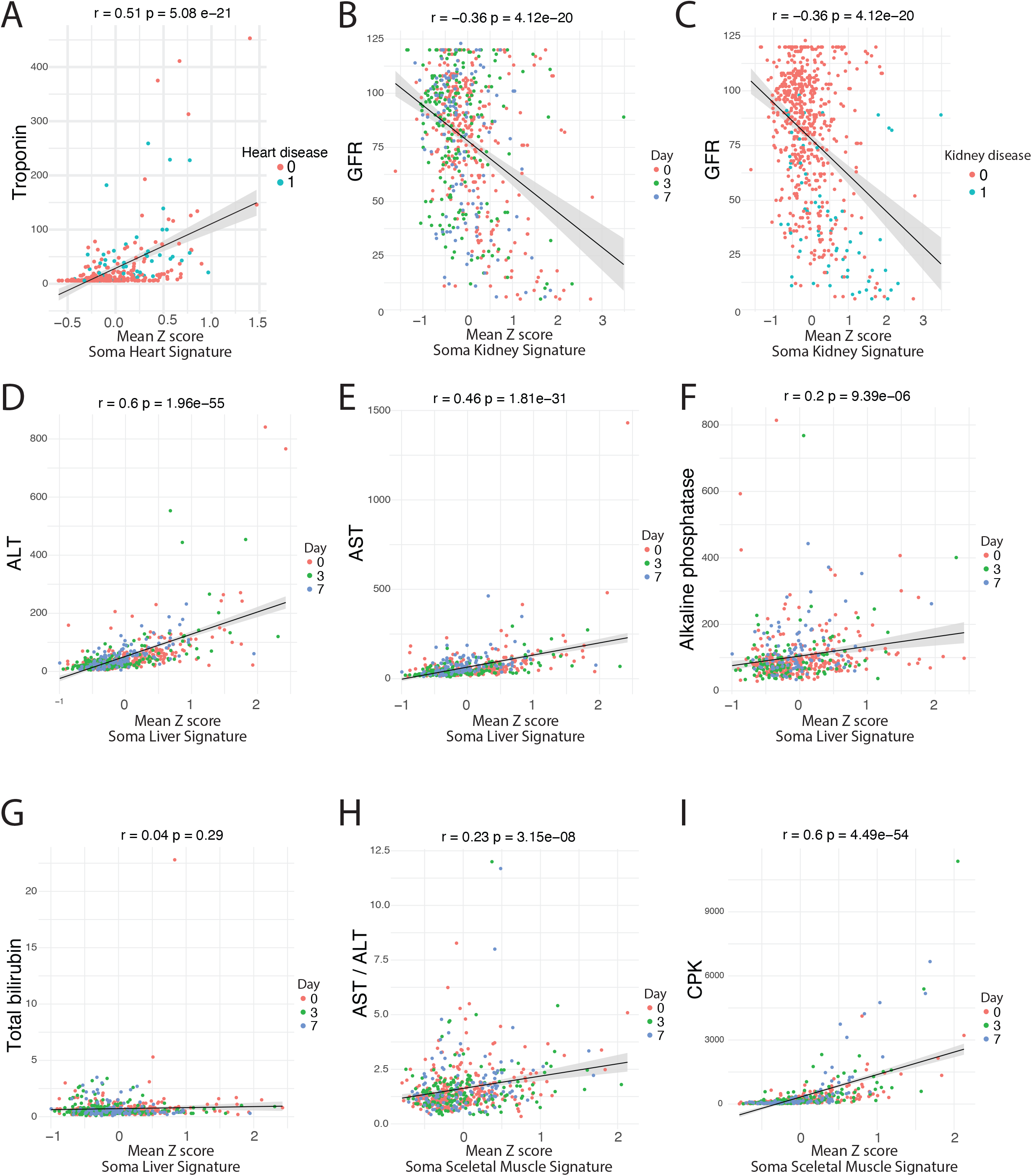
Organ damage plasma protein signature correlations with clinical laboratory values. (A) Correlation of heart-specific plasma protein signature with clinical troponin measurements at day 0. (B)-(C) Correlation of kidney-specific plasma protein signature with glomerular filtration rate (GFR) color-coded by time point of sample collection (B) or previously known kidney disease (C). (D)-(G) Correlation of liver-specific plasma protein signatures with clinical measurements of alanine transaminase (ALT) (D), aspartate transaminase (AST) (E), alkaline phosphatase (F), or total bilirubin (G). (H)-(I) Correlation of skeletal muscle-specific plasma protein signature with AST to ALT ratio (H) or creatinine phosphokinase (CPK) (I).

**Extended Data Figure 17.**
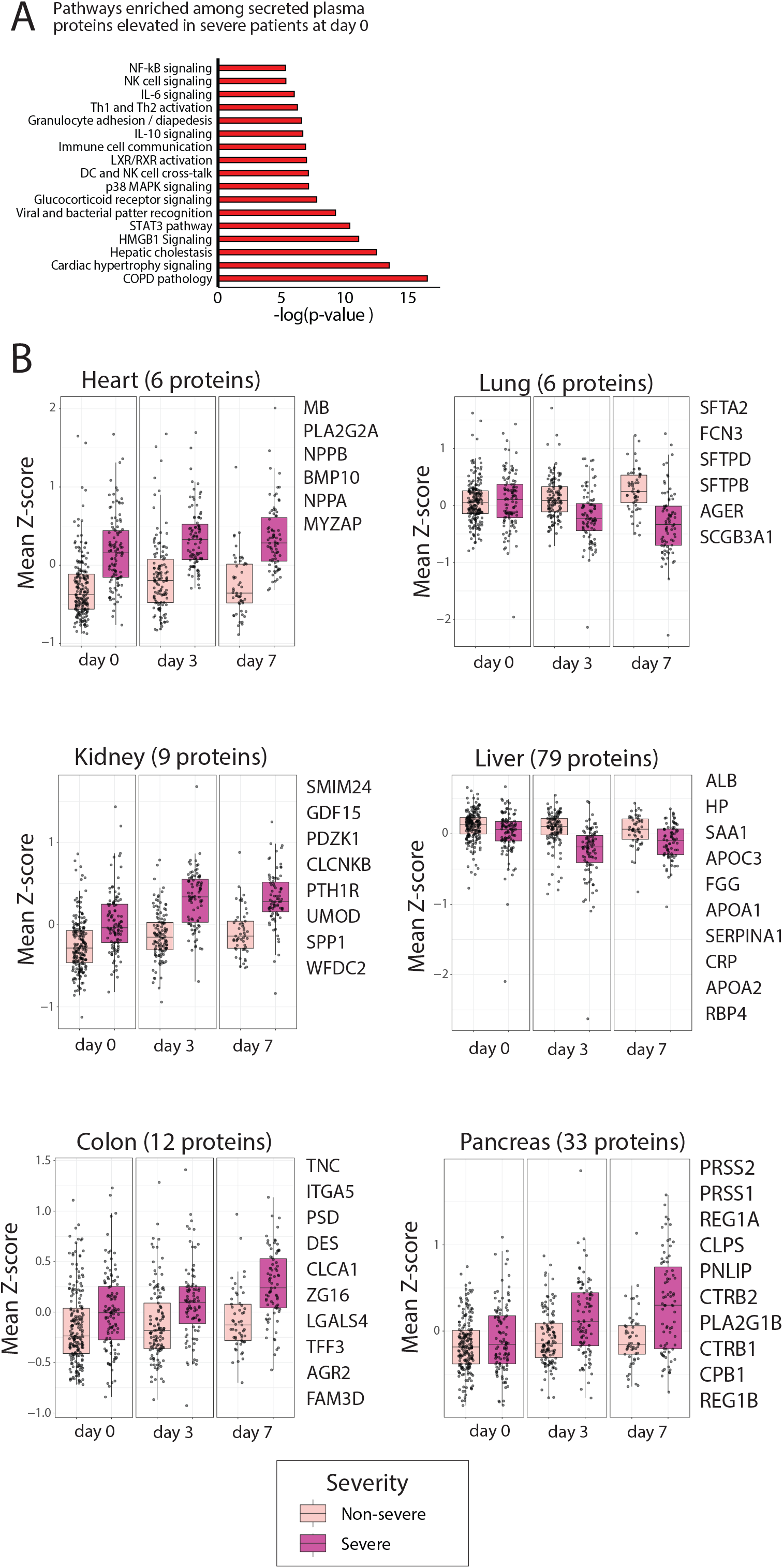
Plasma protein levels of tissue-specific secreted proteins. (A) Pathways enriched amongst plasma proteins that are secreted or membrane-bound and are differentially-expressed at D0 between severe and non-severe COVID-19-positive patients. (B) Expression of derived tissue-specific (heart, lung, kidney, liver, colon, or pancreas) secreted or membrane-bound protein signatures, comparing severe and non-severe patients over time. Shown are the top associated secreted protein with each organ (up to 10).

**Extended Data Figure 18.**
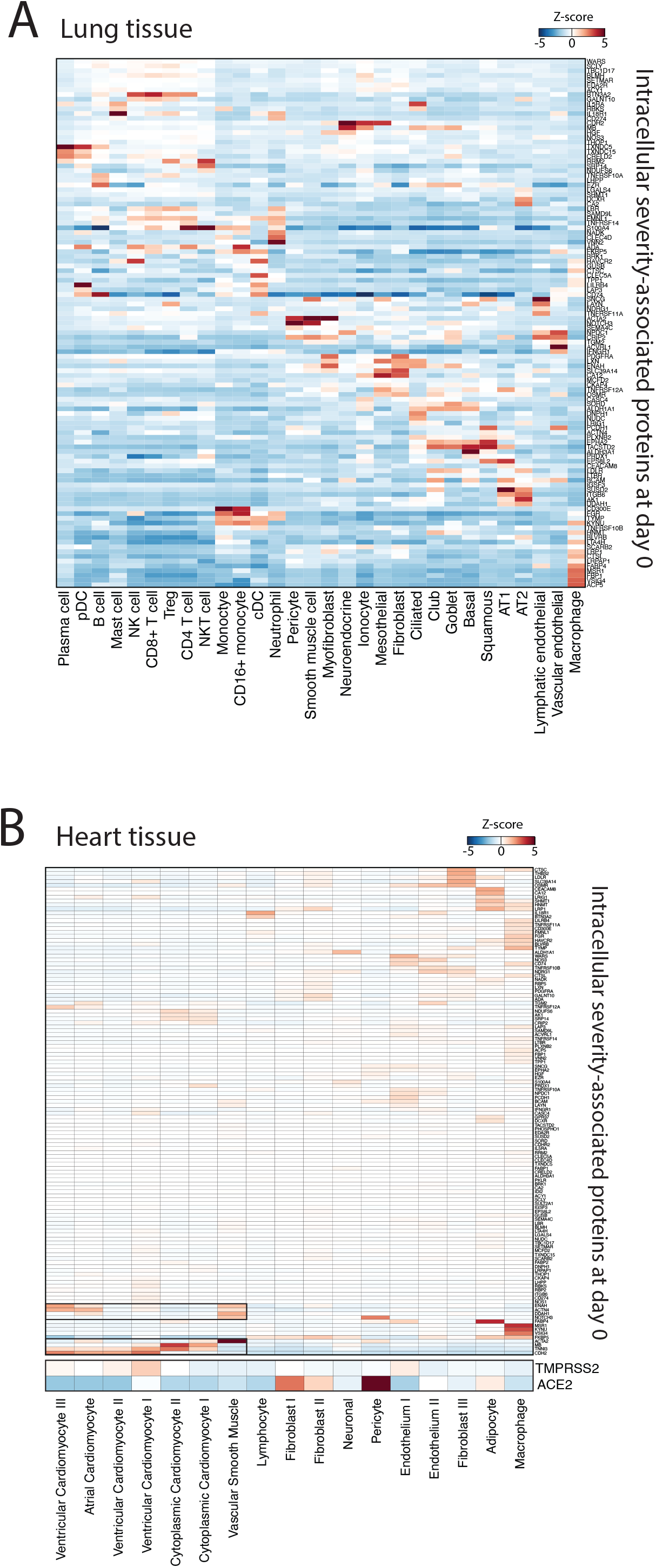
Lung and heart-specific cellular death signatures. (A) Heatmap showing cell-type expression of severity-associated intracellular proteins, derived from SomaScan data mapped to cell types in scRNAseq data of BAL fluid from COVID-19 patients^34^. (B) Heatmap showing cell-type expression of severity-associated intracellular proteins derived from SomaScan data mapped to cell types in scRNAseq data of normal cardiac tissue^53^.

**Extended Data Figure 19.**
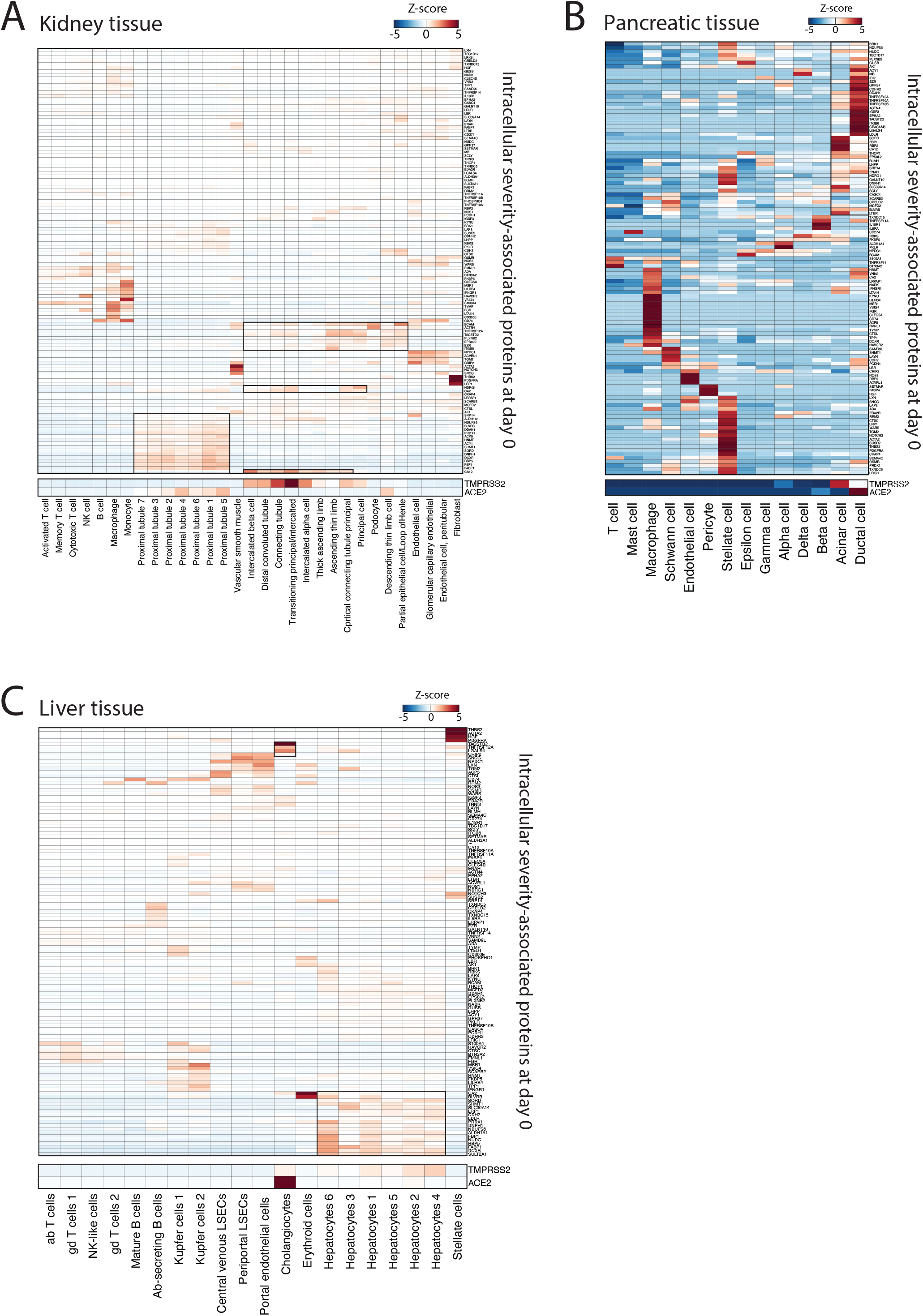
Kidney, pancreatic and liver-specific cellular death signatures. (A) Heatmap showing cell-type expression of severity-associated intracellular proteins derived from SomaScan data mapped to normal kidney tissue^54^. (B) Heatmap showing cell-type expression of severity-associated intracellular proteins derived from SomaScan data mapped to normal pancreatic tissue. (C) Heatmap showing cell-type expression of severity-associated intracellular proteins derived from SomaScan data mapped to normal liver tissue^55^.

**Extended Data Figure 20.**
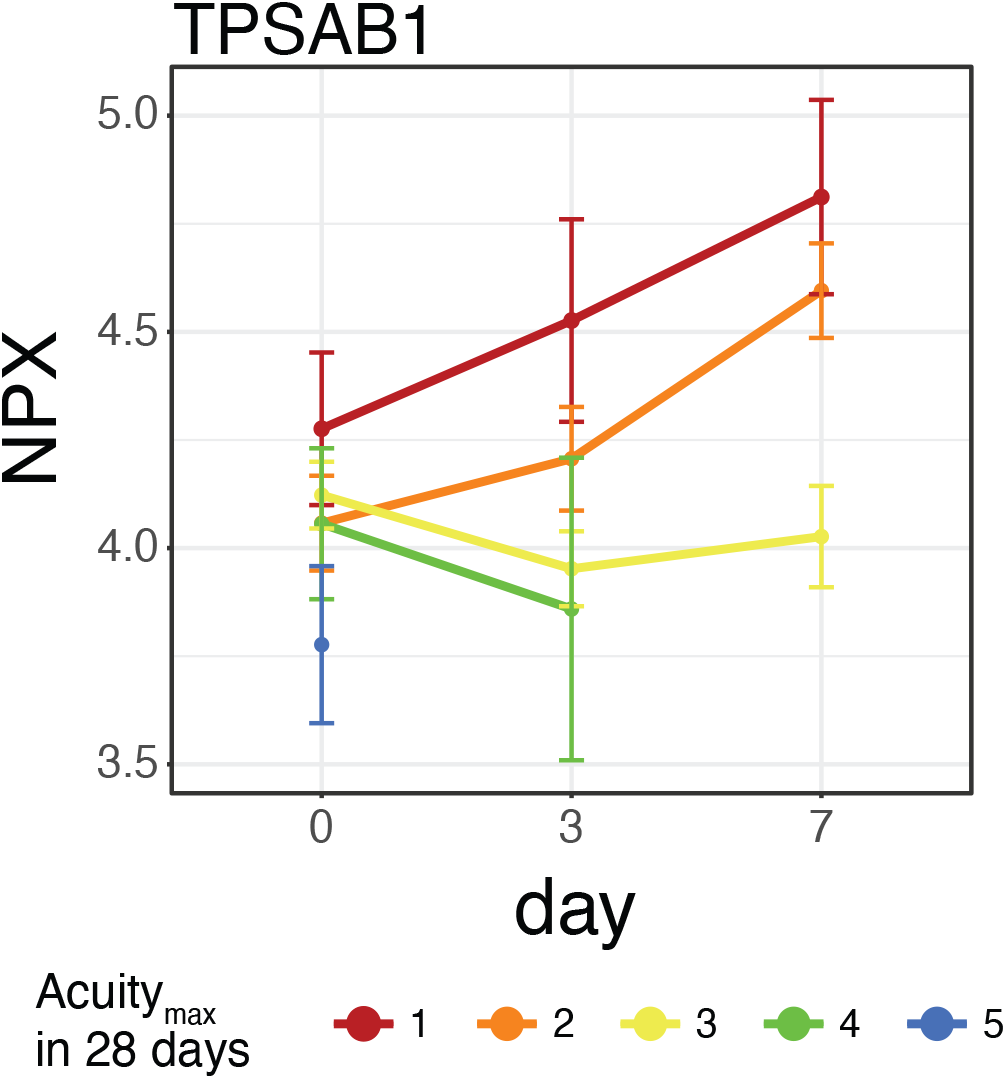
Expression of the mast cell marker tryptase (TPSAB1 assay by Olink) over time, by Acuity_max_ level.

**Extended Data Figure 21.**
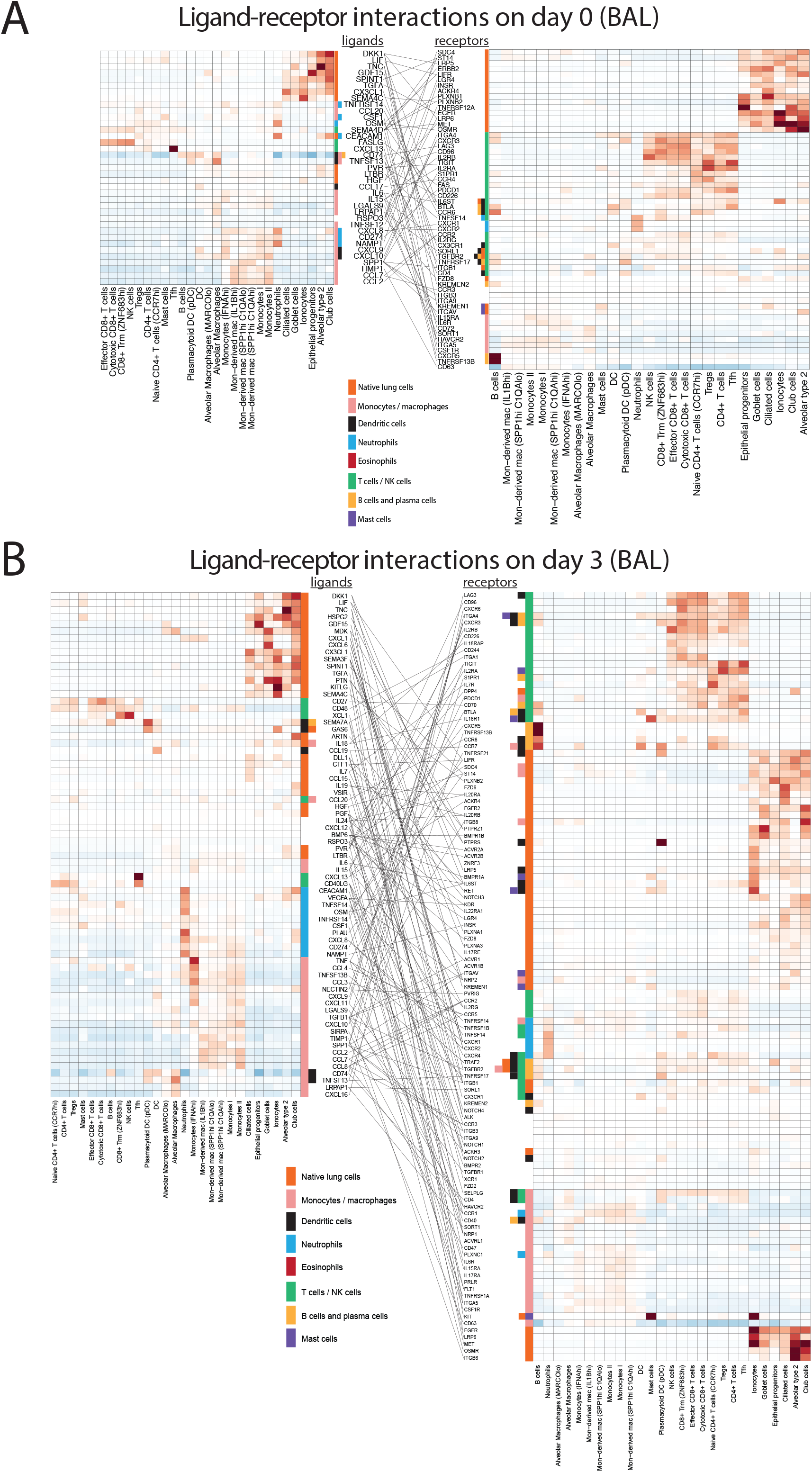
Cellular communication between lung cell subsets in COVID-19-positive patients. (A)-(B) Ligand-receptor relationships of severity-associated plasma ligands shown in BAL fluid of COVID-19^34^ patients for ligands significant on day 0 (A) or on day 3 (B).

## Supplemental tables

**Table S1**. Clinical metadata and associations with outcome for all patients in this cohort.

**Table S2**. List of proteins assayed using the Olink proteomics platform.

**Table S3**. Processed normalized protein expression (NPX) values used for this analysis from the Olink assay.

**Table S4**. Linear model outcome for the comparison of COVID-19-positive and negative patients.

**Table S5**. Linear mixed model outcome for the comparison of severity and time as main effects within COVID-19-positive patients using the Olink assay.

**Table S6**. Linear mixed model outcome for the comparison of Acuity_max_ and time as main effects within COVID-19-positive patients using the Olink assay.

**Table S7**. Linear mixed model outcome for the comparison of Acuity_max_ and time as main effects within COVID-19-positive patients using the Olink assay, with group A1 split into intubated and non-intubated patients.

**Table S8**. Residual values for the linear mixed model outcome using all potentially confounding covariates within COVID-19-positive patients using the Olink assay.

**Table S9**. Clinical characteristics of patient endotypes defined by Louvain clusters.

**Table S10**. Pairwise correlations between patient age and each protein measured using the Olink platform. **Table S11**. Linear mixed model outcome for the comparison of age and time as main effects within COVID-19-positive patients using the Olink assay.

**Table S12**. Linear mixed model outcome for the comparison of age, time and severity as main effects within COVID-19-positive patients using the Olink assay.

**Table S13**. Linear mixed model outcome for the comparison of severity and time as main effects within COVID-19-positive patients using the SomaScan assay.

**Table S14**. Derived organ-specific intracellular plasma protein signatures.

**Table S15**. p-values for survival analysis using organ-specific intracellular plasma protein signatures.

## Acknowledgments

We owe deep gratitude to the participants in this study. We thank the all the clinical nursing staff that made sample collection possible, and the Translational and Clinical Research Center (TCRC) for arranging collection of all follow-up samples on inpatient floors and ICUs, in particular Grace Holland, RN, Katherine Broderick, RN and Siobhan Boyce, RN. We thank the Departments of Emergency Medicine and Medicine, Infection Control and the Center for Disaster Preparedness at Massachusetts General Hospital (MGH) for institutional support to enable enrollment during a time when access to clinical spaces was severely limited. We also thank the Departments of Emergency Medicine and Medicine for their financial support in maintaining needed staffing levels during enrollment, when many research funding sources were being suspended. We thank Caroline Beakes and Nicole Russell for assistance with data entry. Finally, we thank Jayaraj Rajagopal, Itai Yanai, Patrick Ellinor and Mark Chaffin for access to their processed single-cell RNA-sequencing datasets.

Direct funding for this project was provided in part by a grant from the National Institute of Health (N.H., U19 AI082630), an American Lung Association COVID-19 Action Initiative grant (M.B.G.), and grants from the Executive Committee on Research at MGH (M.B.G.), the Chan-Zuckerberg Initiative (A-C.V.). N.H. was also funded by a gift from Arthur, Sandra and Sarah Irving for the David P. Ryan, MD Endowed Chair in Cancer Research. This work was also supported by the Harvard Catalyst / Harvard Clinical and Translational Science Center (National Center for Advancing Translational Sciences, National Institutes of Health Awards UL1 TR 001102 and UL1 TR 002541-01). We are also very grateful for the generous contributions of Olink Proteomics Inc. and Novartis (in collaboration with SomaLogic,Inc.) for providing in-kind all proteomics assays presented in this work, without which our findings would not have been possible.

## Author information

These authors jointly supervised this work: Michael R. Filbin, Arnav Mehta, Nir Hacohen, Marcia B. Goldberg.

## Author Contributions

Conceptualization: Filbin (M.R.F.), Mehta (A.M.), Hacohen (N.H.), Goldberg (M.B.G.);

Resources: Filbin (M.R.F.), Hacohen (N.H.), Goldberg (M.B.G.), Grundberg (I.G.), and Jennings (L.L.J);

Methodology: Mehta (A.M.), Hacohen (N.H.), Filbin (M.R.F.), Goldberg (M.B.G.), Sade-Feldman (M.S-F.), Villani (A-C.V.), Parry (B.A.P), Bhattacharyya (R.P.B.), Jennings (L.L.J.), Grundberg (I.G.), and Gerszten (R.E.G.);

Investigation: (All) M.R.F., A.M., A.M.S., K.R.K., J.R.G., M.G., B.G.F., N.C.C., A.L.K.G., I.G., H.K.K., T.J.L., K.M.L-P., B.M.L., C.L.L., K.M., J.D.M., B.N.M., M.R-L., B.C.R., N.S., J.T., M.F.T., R.E.G., G.S.H., P.J.H., D.J.L., B.L., D.N., K.P., M.R., C.S.S., A.W., T.E.W., A.S.Z., L.L.J., I.G., R.P.B., B.A.P., A-C.V., M.S-F., N.H., and M.B.G.;

Formal Analysis: Mehta (A.M.), Schneider (A.M.S.), Guess (J.R.G.), Gentil (M.G.), Fenyves (B.G.F.) and Lin (B.L.).

Writing – Original Draft: Mehta (A.M.), Filbin (M.R.F.), Hacohen (N.H.), and Goldberg (M.B.G.);

Writing – Review & Editing: Filbin (M.R.F.), Mehta (A.M.), Hacohen (N.H.), Goldberg (M.B.G.), Schneider (A.M.S.), Gerstzen (R.E.G.), Fenyves (B.G.F.), Grundberg (I.D.), Jennings (L.L.J.), Villani (A-C.V.), Sade-Feldman (M.S-F.), Zajak (A.S.Z.), Wood (T.E.W.), Russo (B.R.), Bhattacharyya (R.P.B.), and Parry (B.A.P.).

## Corresponding authors

Correspondence to Marcia B. Goldberg, Nir Hacohen, Michael Filbin, Arnav Mehta.

## Declaration of Interests

A.M.: Consultant for Third Rock Ventures

J.R.G. & I.G.: Employee of Olink Proteomics

G.S.H.: Employee of Genentech (as of November 2020)

L.L.J.: Employee and stockholder of Novartis

N.H.: Holds equity in BioNTech and is a consultant for Related Sciences

## Funding Sources

M.R.F.: Research grants received from Day Zero Diagnostics, Rapid Pathogen Screening Inc., Nihon Kohden Corporation (all unrelated to this work)

A.M.: NIH / NCI T32 2T32CA071345-21A1 (unrelated to this work)

M.G.: Recipient of an EMBO Long-Term Fellowship (ALTF 486-2018) and a Cancer Research Institute / Bristol-Myers Squibb Fellow (CRI2993) (unrelated to this work)

B.G.F.: Rosztoczy Foundation Scholarship (unrelated to this work)

A-C.V.: NIH / NCI DP2CA247831 and Damon Runyon-Rachleff Innovation Award (unrelated to this work)

G.S.H.: James S. McDonnell Foundation Postdoctoral Fellowship (unrelated to this work)

B.L.: Cystic Fibrosis Foundation Postdoctoral Fellowship, LIN19F0

A.S.Z.: NIAID F32 AI145128-01A1 (unrelated to this work)

B.A.P.: Supported by 1U24NS100659-01 & 1U10NS080369-01 (unrelated to this work)

N.H.: NIH / NIAID U19 AI082630; Chair and gift from Sandra, Sarah and Arthur Irving (unrelated to this work)

M.B.G.: Research grants from American Lung Association (COVID-19 Action Initiative), Executive Committee on Research at MGH

